# The role of insulators and transcription in 3D chromatin organisation of flies

**DOI:** 10.1101/2021.04.26.441424

**Authors:** Keerthi T Chathoth, Liudmila A Mikheeva, Gilles Crevel, Jareth C. Wolfe, Ioni Hunter, Saskia Beckett-Doyle, Sue Cotterill, Hongsheng Dai, Andrew Harrison, Nicolae Radu Zabet

## Abstract

The DNA in many organisms, including humans, is shown to be organised in topologically associating domains (TADs). In *Drosophila*, several architectural proteins are enriched at TAD borders, but it is still unclear whether these proteins play a functional role in the formation and maintenance of TADs. Here, we show that depletion of BEAF-32, Cp190, Chro and Dref leads to changes in TAD organisation and chromatin loops. Their depletion predominantly affects TAD borders located in heterochromatin, while TAD borders located in euchromatin are resilient to these mutants. Furthermore, transcriptomic data has revealed hundreds of genes displaying differential expression in these mutants and showed that the majority of differentially expressed genes are located within reorganised TADs. Our work identifies a novel and functional role for architectural proteins at TAD borders in *Drosophila* and a link between TAD reorganisation and subsequent changes in gene expression.

## Introduction

Topologically associating domains (TADs) provide a fundamental unit for chromosome organisation^1, 2^ and are widely conserved across species^3^ as well as during different developmental stages^4, 5^, suggesting that they have a functional role. Furthermore, in *Drosophila* cells, changes in the 3D organisation of DNA after heat stress have been found to correlate with transcriptional changes^6^. Recent evidence points to defective 3D architecture as a major contributor for diseases, developmental defects and even ageing^7–13^. These results suggest that 3D organisation of the DNA is important in gene regulation.

There has been significant progress in generating empirical data on chromatin organisation in different organisms and tissues, but, despite this, the mechanisms that drive the formation of TAD borders remain unclear. Previous research has shown that TAD borders are enriched in housekeeping genes^6^, developmental enhancers^14^ and highly conserved genomic regulatory blocks^15^. In addition, architectural proteins and insulators are enriched at TAD borders^16, 17^. Two different mechanisms were proposed to be responsible for TAD formation: *(i)* compartment domains, which are formed by interactions among sequences that contain active or inactive histone modifications and *(ii)* loop domains that are flanked by CTCF binding sites and are formed by a cohesin driven loop extrusion mechanism^18–20^. The latter displays a strong loop localised at the top of the TAD, while the former lacks this chromatin loop. In mammalian systems, CTCF and cohesin are the main architectural components that are located at TAD borders and their depletion has been shown to disrupt TADs^21–23^. By contrast, in *Drosophila*, several insulator proteins occupy TAD borders, such as CTCF, BEAF-32, Chro and Cp190^16, 24–27^, but the majority of TADs lack the chromatin loop at the top of the TAD suggesting a prevalence of the compartment domains^27, 28^. In particular, previous research has identified strong enrichment of BEAF-32 at TAD borders in *Drosophila*^16, 25, 26, 29^, but this was more pronounced in cell lines derived from the embryo (Kc167 derived from dorsal closure stage and S2 derived from late embryonic stage) or whole embryos. Interestingly, there are negligible changes in 3D chromatin organisation following BEAF-32 RNAi knockdown in Kc167 cells^25^ despite the strong enrichment of BEAF-32 at TAD borders. Kc167 cells display saturating levels of BEAF-32 at TAD borders, suggesting that upon RNAi knockdown, there is potentially still sufficient protein present in the cell to maintain TAD borders^30^. Furthermore, BEAF-32 displays the same binding motif as another architectural protein in *Drosophila* called Dref^31, 32^. When BEAF-32 is depleted, one possibility is that Dref replaces it at TAD borders and this could explain the lack of changes in 3D organisation observed in Kc167 cells.

Two additional proteins, Cp190 and Chro, are enriched at TAD borders^14, 24, 29^. These proteins cannot bind independently to DNA, but are recruited mainly by BEAF-32^33^, with up to 91% of TAD borders in a *Drosophila* cell line (S2) displaying presence of BEAF-32 together with either Cp190 or Chro^29^. Like BEAF-32, the role of Cp190 and Chro at TAD borders is currently unclear.

Recently, the role of TADs in gene regulation has been challenged ^34, 35^. In one example, it was shown that changes in TAD borders and changes in transcription are not coupled when investigating a *Drosophila* balancer chromosome containing chromosome re-arrangements^34^. However, the balancer chromosomes display a very small number of rearrangements that result in changes at only a few TAD borders. It is less likely that effects on gene expression will be observed when sampling only a few re-arrangements and one possibility is that more and stronger changes in TADs (e.g., more TAD borders are lost) would allow the observation of changes in gene expression that correlate with reorganisations of TADs.

We depleted BEAF-32 in BG3 cells (derived from the larval central nervous system) using RNAi knockdown and measured the changes in 3D chromatin organisation at sub-kilobase resolution together with changes in transcription to dissect the mechanism at TAD borders and evaluate the functional role of TADs. In BG3 cells, BEAF-32 has reduced levels at TAD borders^26^, which raises the question of whether a strong depletion combined with the low levels of BEAF-32 is sufficient to affect the borders of the TADs. We also performed double knockdowns of Cp190/Chro and BEAF-32/Dref using RNAi to disentangle the interactions between different architectural proteins at TAD borders.

## Results

### BEAF-32, Cp190 and Chro have a functional role at TAD borders in BG3 cells

We performed single knockdown of BEAF-32 and combinatorial knockdown of Cp190 and Chro in BG3 cells followed by *in situ* Hi-C (Figure S1 and Tables S1 and S2). The RNAi knockdowns lead to specific and strong reduction in both the mRNA levels and protein levels and do not affect the cell cycle (Figure S1). High-resolution contact maps were generated for both knockdown mutants. The biological replicates displayed high similarities and were merged for the downstream analysis (Figure S2). There was a noticeable re-organisation in the contact maps as a result of the knockdowns when compared to wild type BG3 cells (Figure S2 and Figure 1A). BG3_BEAF-32_^-^ resulted in loss of long-range interactions and showed an increase in short-range interactions (Figure 1A). Likewise, BG3_Cp190_^-^_Chro_^-^ also exhibited reduced long-range interactions and increased short-range interactions, but the loss of long-range interactions were less pronounced compared to BG3_BEAF-32_^-^ (Figure 1A).

**Figure 1.**
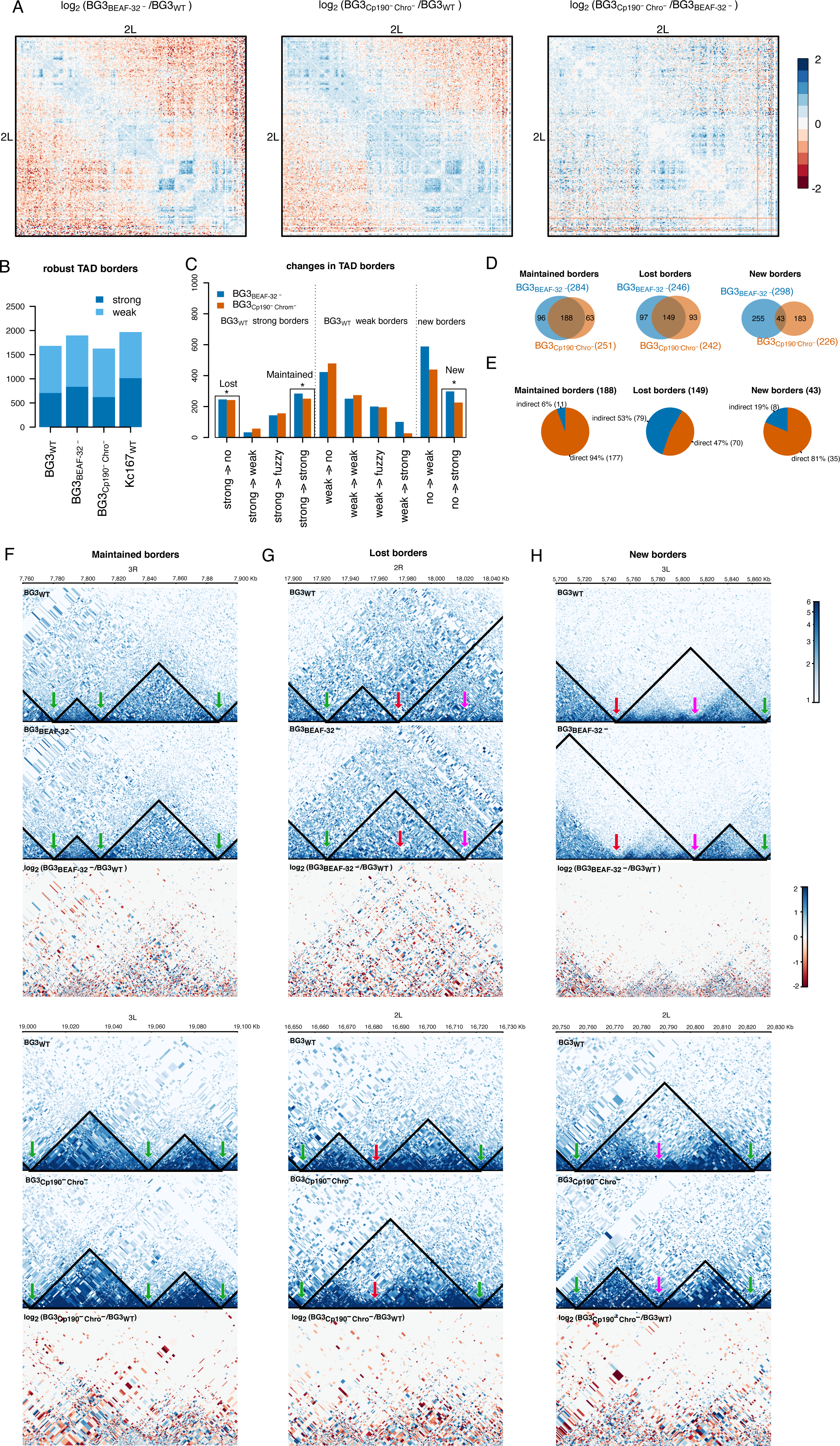
Functional role of BEAF-32, Cp190 and Chro in TAD organisation of BG3 cells. (A) log_2_ ratio between the normalised number of contacts on chromosome 2L in: *(i)* BEAF-32 knockdown and WT cells, *(ii)* Cp190 Chro double knockdown and WT cells and *(iii)* Cp190 Chro double knockdown and BEAF-32 knockdown cells. Red colours indicate less contacts in the first condition and blue colours more contacts in the first condition. (B) Number of robust TAD borders in BG3 cells (WT, BEAF-32 knockdown and Cp190 Chro double knockdown) (see Figure S3). We also included the number of TAD borders in Kc167 cells. We split each class of TAD border into two subgroups: strong borders and weak borders, depending on whether the TAD borders can still be detected when increasing the stringency of the TAD calling algorithm. (C) Classification of TAD borders as described in the main text: lost (borders that are strong in WT and completely disappear in the mutant), maintained (borders that are strong in WT and are maintained strong in the mutant) and new (borders that appear strong in the mutant). (D) Overlap of lost, maintained and new borders in the two mutants. (E) Number and percentage of maintained, lost and new borders that have direct binding of BEAF-32, Cp190 and/or Chro (see Figure S4A). We considered common borders between the two mutants (BEAF-32 single knockdown and Cp190 and Chro double knockdown). (F-H) Examples of genomic regions at DpnII restriction size resolution for: (F) maintained, (G) lost and (H) new TAD borders. Darker colours indicate more contacts retrieved by *in situ* Hi-C while the black line indicates the TADs. Green arrows indicate maintained borders, red arrows lost borders and purple arrows new borders. From top to bottom we plot the contact map and TADs in WT cells, in the mutants (BEAF-32 knockdown and Cp190 Chro double knockdown) and log_2_ of the ration between mutant and WT.

Several papers have proposed that BEAF-32, Cp190 and Chro control the borders of TADs ^16, 25, 29^. We then investigated the TAD border classification as performed previously^26^. In particular, we used HiCExplorer^25^ and identified between 2000 and 2600 TADs (see Table S2 and Methods), which is consistent with other studies^14, 25, 26^. TAD borders were classified into weak and strong borders depending on whether they can be detected with increasing stringency of the TAD calling algorithm, with strong borders being detected even with the more stringent parameters (see Methods). To investigate the robustness of these TAD borders, we downsampled all Hi-C libraries by 20% and repeated the analysis (see Figure S3). 706 of the 989 strong TAD borders in WT cells are robust, meaning they are recovered in both full and downsampled datasets (Figure 1B).

Compared to WT BG3 cells, out of all strong borders (706), 188 borders were maintained and 149 were lost in both mutants (BG3_BEAF-32_^-^ and BG3_Cp190_^-^_Chro_^-^) (see Figure 1C-D and Methods). In both knockdowns approximately 150 strong borders and 200 weak borders changed their position within 2Kb and we called them fuzzy borders. In addition, approximately one quarter of 975 weak borders from BG3 WT cells were maintained as weak borders in the two mutants, but only a negligible number of borders converted from strong to weak or vice versa (Figure 1C).

Next, in order to distinguish between direct and indirect effects, we evaluated how many of the maintained and lost robust borders have BEAF-32, Chro or Cp190 ChIP peaks in their vicinity. Figure 1E shows that majority of maintained TAD borders (94%) are direct targets of the three proteins, but only half of the lost TAD borders (47%) are direct targets (also see Figure S4A). Furthermore, the majority of maintained TAD borders (70%) retain BEAF-32 or Cp190 upon knockdown, but most of the lost borders (70%) lose binding of these architectural proteins after knockdown (Figure S4B-C). This further confirms that the direct maintained and direct lost TAD borders are indeed controlled by the three architectural proteins.

Some regions displayed high conservation of the TAD structure organisation (Figure 1F), while others showed reorganisation (Figure 1G-H). We observed that a loss of a TAD border could result in either movement of the TAD borders (see top panel in Figure 1G) or aggregation of two TADs (see bottom panel in Figure 1G).

We also found new border formation in both knockdowns, ranging between 400 to 600 weak borders and 200 to 300 strong borders. The majority of these new borders moved more than 2Kb in the mutants compared to WT (Figure 1G-H and Figure S5). A small proportion of the new TAD borders result in splitting the original TAD in two separate TADs (see bottom panel of Figure 1H and Figure S5). Out of all the new borders, only 43 were common between both knockdowns (Figure 1D). This may be explained by the fact that Chro and Cp190 are able to bind chromatin independent of BEAF-32^36^. Interestingly, the majority of these new borders have BEAF-32, Chro or Cp190 ChIP peaks in their vicinity and retain BEAF-32 or Cp190 upon knockdown (Figure 1E and Figure S4A-C). To identify the roles of BEAF-32, Cp190 and Chro at TAD borders, we focused on two groups: *(i)* maintained borders (robust TAD borders that are strong in WT cells and are maintained strong in both mutants) and *(ii)* lost borders (robust TAD borders that are strong in WT cells and are lost in the two mutants).

### Combined Dref and BEAF knockdown shows an enhanced effect on TAD border distribution

Dref is a DNA binding protein that shares a similar binding motif with BEAF-32, meaning that upon depletion of BEAF-32, Dref could potentially replace it at TAD borders. To investigate this, we performed a combinatorial knockdown of BEAF-32 and Dref (Figure S1) followed by *in situ* Hi-C (Tables S1 and S2). Again, the combinatorial knockdown resulted in specific and efficient depletion at both mRNA and protein levels and does not affect the cell cycle (Figure S1). In the BEAF-32 Dref double knockdown (BG3_BEAF-32_^-^_Dref_^-^) we noticed a more pronounced effect in the reorganisation of the 3D interaction compared to the BG3_BEAF-32_^-^ or BG3_Cp190_^-^_Chro_^-^ mutants (Figure 2A). In particular, BG3_BEAF-32_^-^_Dref_^-^ displayed significantly fewer robust TAD borders (982) of which only one third are strong (292) (Figure S3); with the majority of the TAD borders being lost (Figure 2B). There were 50% more TAD borders that were lost in BG3_BEAF-32_^-^_Dref_^-^ compared to the single knockdown of BEAF-32 or double knockdown of Cp190 and Chro (Figure 2C, D and F). While looking at the maintained borders, only one third of the borders were maintained in BG3_BEAF-32_^-^_Dref_^-^ when compared to BG3_BEAF-32_^-^ or BG3_Cp190_^-^_Chro_^-^ (Figure 2C, D and E). In addition, 161 new borders appear in the BG3_BEAF-32_^-^_Dref_^-^ double knockdown (see Figure 2C and D). The majority of these new borders are movements of borders in the mutant compared to the closest WT border (Figure S5 and Figure 2G). Overall, we found that there is a large overlap between TAD borders that are lost in the three mutants and also a large subset of TAD borders that disappear only in the BG3_BEAF-32_^-^_Dref_^-^ mutant, indicating that there is a subset of TAD borders that require Dref for maintenance (Figure 2C).

**Figure 2.**
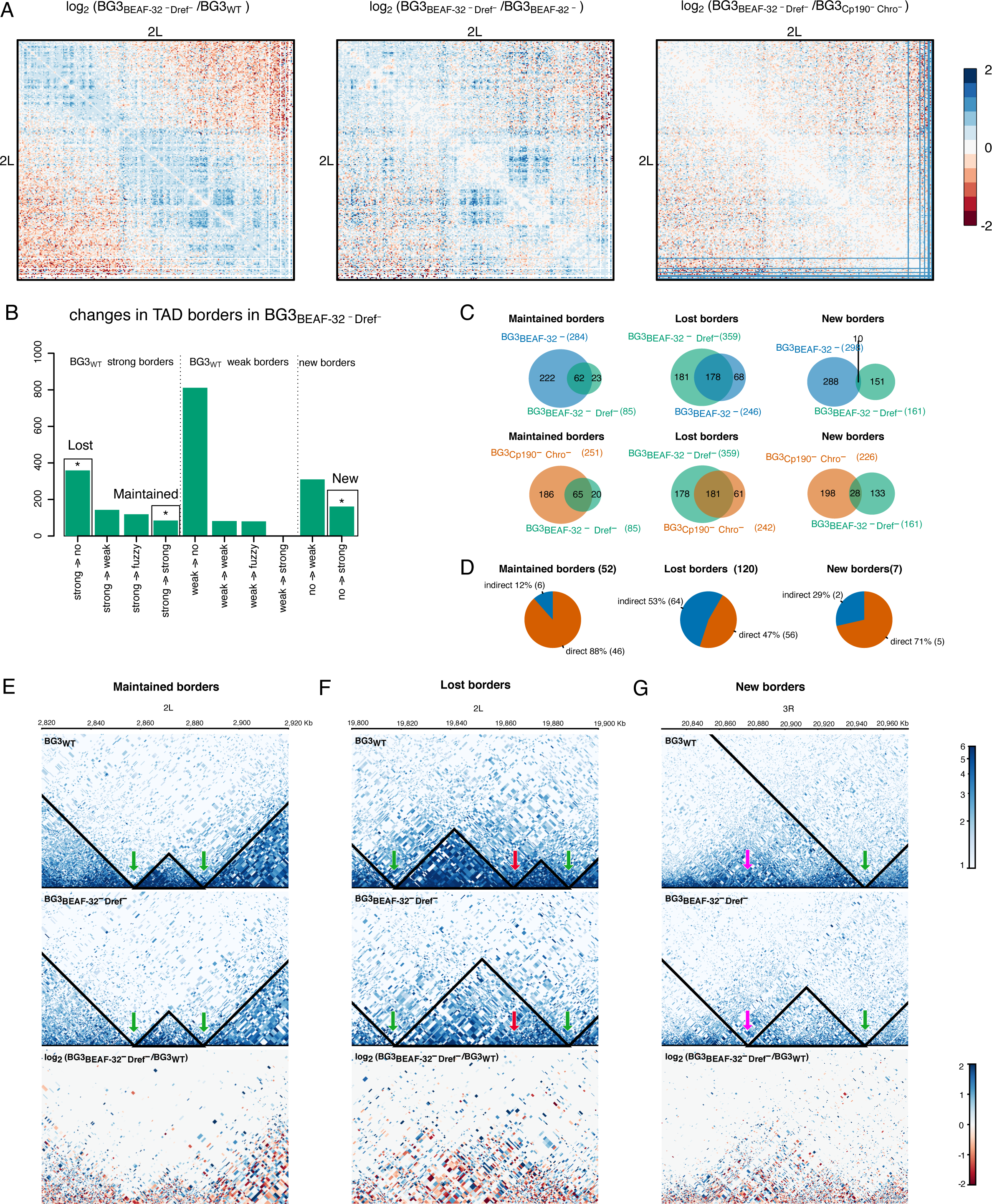
Functional role of Dref in 3D chromatin organisation of BG3 cells. (A) log_2_ ratio between the normalised number of contacts on chromosome 2L in: *(i)* BEAF-32 Dref double knockdown and WT cells, *(ii)* BEAF-32 Dref double knockdown and BEAF-32 knockdown cells and *(iii)* BEAF-32 Dref double knockdown and Cp190 Chro double knockdown cells. Red colours indicate less contacts in the first condition and blue colours more contacts in the first condition. (B) Classification of robust TAD borders as described in the main text: lost (borders that are strong in WT and completely disappear in the mutant), maintained (borders that are strong in WT and are maintained strong in the mutant) and new (borders that appear strong in the mutant); see Figure S3. (C) Overlap of lost, maintained and new borders in the three mutants. (D) Number and percentage of maintained, lost and new borders that have direct binding of BEAF-32, Cp190 and/or Chro (see Figure S4B). We considered common borders between the all three mutants (BEAF-32 single knockdown, Cp190 and Chro double knockdown and BEAF-32 and Dref double knockdown). (E-G) Examples of genomic regions at DpnII restriction size resolution for: (E) maintained, (F) lost and (G) new TAD borders. Darker colours indicate more contacts retrieved by *in situ* Hi-C while the black line indicates the TADs. Green arrows indicate maintained borders, red arrows lost borders and purple arrows new borders. From top to bottom we plot the contact map and TADs in WT cells, in the mutants (BEAF-32 Dref double knockdown) and log_2_ of the ration between mutant and WT.

To distinguish the direct targets from indirect, we aligned TAD borders with the protein occupancy (see methods and Figure S4D). The majority of TAD borders that are maintained in BG3_BEAF-32_^-^_Dref_^-^ (and also in BG3_BEAF-32_^-^ and BG3_Cp190_^-^_Chro_^-^) (88%) are direct targets of BEAF-32, Cp190 and Chro (see Figures 2D and S4D). However, only half of the lost TAD borders in BG3_BEAF-32_^-^_Dref_^-^ (47%) (and also in BG3_BEAF-32_^-^ and BG3_Cp190_^-^_Chro_^-^) are direct targets of the three proteins. Upon knockdown of BEAF-32 or Cp190, the majority of the maintained TAD borders in BG3_BEAF-32_^-^_Dref_^-^

(70%) retain BEAF-32 or Cp190 and most of the lost borders in BG3_BEAF-32_^-^_Dref_^-^ (65%) have lost occupancy of these proteins (Figure S4E-F). Similar to maintained borders, the majority of new borders display binding of BEAF-32, Chro and/or Cp190. Binding of BEAF-32 or Cp190 is retained at these new borders upon knockdown. These results are similar to the ones for the maintained, lost and new borders common between BG3_BEAF-32_^-^ and BG3_Cp190_^-^_Chro_^-^

Interestingly, the majority of TAD borders that are lost only in BG3_BEAF-32_^-^_Dref_^-^ (and are maintained in BG3_BEAF-32_^-^ or BG3_Cp190_^-^_Chro_^-^) are bound by BEAF-32, Cp190 and/or Chro in WT cells (Figure S4G-H). This suggests that Dref displays redundancy to BEAF-32, by maintaining TAD borders when BEAF-32 is absent. When both architectural proteins are depleted then these TAD borders that were maintained after BEAF-32 single knockdown are also lost.

### Reorganisation in TADs correlates with changes in gene expression

Several studies have shown that TAD reorganisation leads to changes in transcription that correspond to developmental defects or diseases^7–12^. Nevertheless, other studies failed to find a connection between changes in TADs and transcription^34, 35^. Here, instead of disrupting TADs by rearrangements of the DNA at TAD borders, we perturbed a large number of TADs by knocking down architectural proteins and investigated whether that leads to changes in gene expression. We found significant changes in gene expression with 598, 688 and 814 differentially expressed genes (DEG) in BG3_BEAF-32_^-^, BG3_Cp190_^-^_Chro_^-^ and BG3_BEAF-32_^-^_Dref_^-^ respectively (Figure 3 and Table S3). The majority of DEGs are upregulated in the mutants compared to WT. Interestingly, almost all of these are found inside robust TADs in both WT and mutants. Figure 3 shows that very few DEGs are in TADs that have both borders conserved in the mutants (10.9% in BG3_BEAF-32_^-^, 8.7% in BG3_Cp190_^-^_Chro_^-^ and 0.1% in BG3_BEAF-32_^-^_Dref_^-^). This means that majority of DEGs (at least 89%) are located in TADs where at least one of the borders moves in the mutants. There was a large number of DEGs in BG3_BEAF-32_^-^_Dref_^-^ where both TAD borders are lost (or move more than 2 Kb away), but this could be a consequence of the reduced number of TADs in that mutant and the corresponding loss of TAD borders. Our results showed that the association of the DEG with reorganised TADs is statistically significant for BG3_BEAF-32_^-^ and BG3_BEAF-32_^-^_Dref_^-^ mutants, specifically, for TADs in which the borders move more than 2 Kb away (Figure S6B). While there are many DEGs in BG3_Cp190_^-^_Chro_^-^, their association with reorganised TADs is not significant. These changes in border positioning cover several massive rearrangement scenarios, such as significant disruption of WT TADs, aggregation of several WT TADs or combination of both. DEGs are randomly distributed inside TADs (no gene spanning over multiple TADs; Figure 3) with no specific localisation near or away from TAD borders (see Figure S7). Our results show that mainly large reorganisations of TADs correspond to significant changes in gene expression and explain why previous studies found contradicting results when establishing a link between TADs and gene expression.

**Figure 3.**
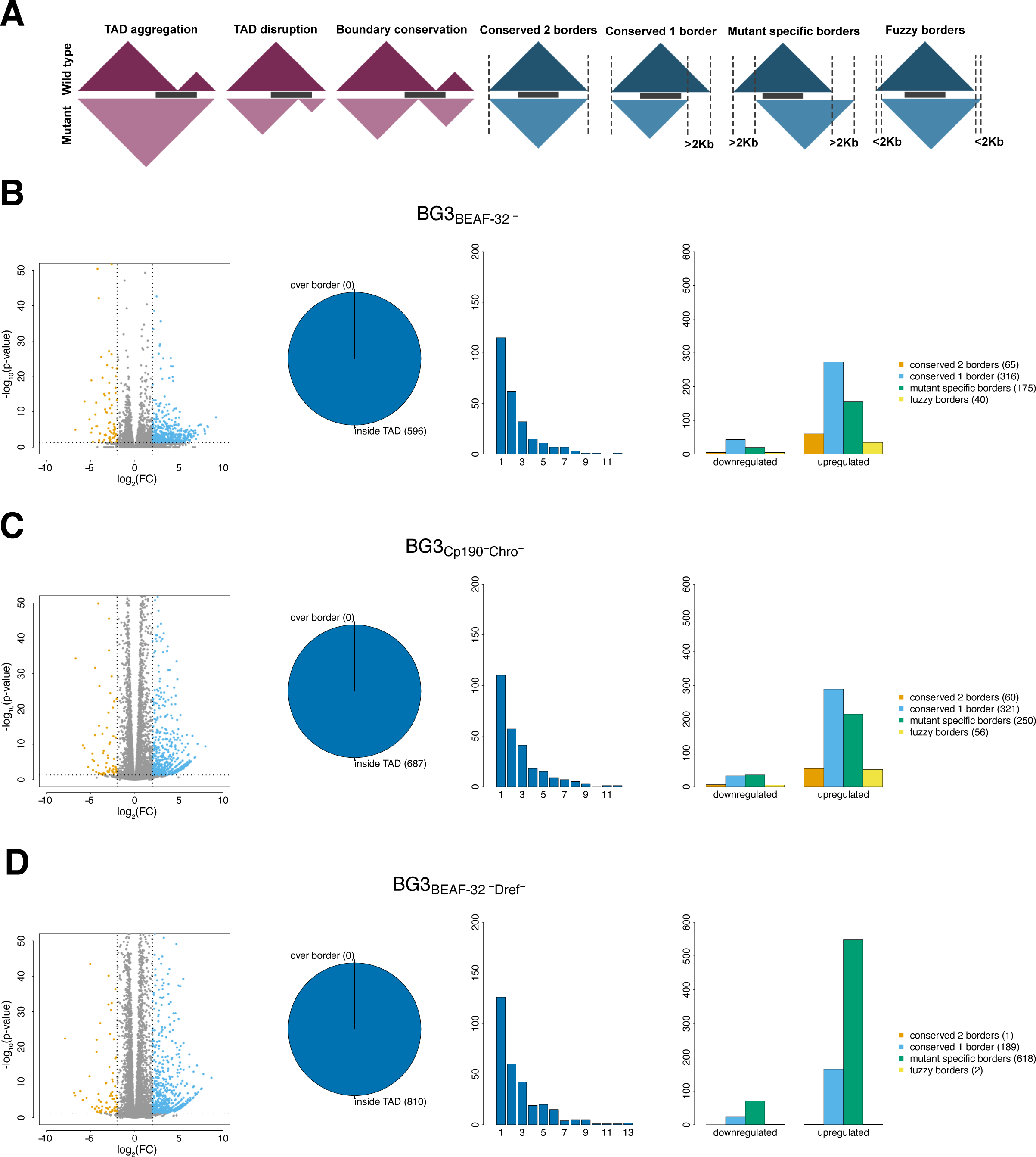
The effects of TAD reorganisation on transcription. (A) The different cases for position of genes in TADs and how the TADs change in the mutant (red for cases where the gene spans over TAD borders and blue for the cases where the gene is within the TADs). (B-D) From left to right: *(i)* volcano plot (orange represents downregulated genes, blue upregulated and grey non-DEG), *(ii)* pie chart with number of differentially expressed genes (over boarder represents red scenario from A and inside TADs represents blue scenario from A), *(iii)* histogram with the number of DEGs in TADs (large number of TADs have more than one DEG) and *(iv)* barplot with the number of downregulated and upregulated genes in different cases where the gene is within the TADs (orange – both TAD borders are conserved, blue – only one of the TAD border is conserved, green – none of the TAD border is conserved and yellow – TAD borders are shifted within 2 Kb). We plot: (B) BEAF-32 knockdown, (C) Cp190 Chro double knockdown and (D) BEAF-32 Dref double knockdown.

### TAD borders are maintained by architectural proteins, divergent transcription and associated factors

In BG3_BEAF-32_^-^ and BG3_Cp190_^-^_Chro_^-^ mutants, we identified two classes of TAD borders: *(i)* maintained in both mutants and *(ii)* lost in both mutants. Given that very few TAD borders are maintained in BG3_BEAF-32_^-^_Dref_^-^, while the majority are lost, we did not include this in the downstream analysis; i.e., the majority of TAD borders that are lost in BG3_BEAF-32_^-^ and BG3_Cp190_^-^_Chro_^-^ are also lost in BG3_BEAF-32_^-^_Dref_^-^, but only a few that are maintained in BG3_BEAF-32_^-^ and BG3_Cp190_^-^_Chro_^-^ are also maintained in BG3_BEAF-32_^-^_Dref_^-^. Furthermore, we selected maintained and lost TAD borders that display binding in WT cells of BEAF-32, Cp190 or Chro and classified these as direct maintained and lost borders.

To determine the chromatin and epigenetic mechanisms present at maintained and lost borders, we analysed the presence of key factors (such as architectural proteins, transcription, replication and accessibility related complexes) at direct lost and direct maintained borders after knockdown. CTCF was partially present at the maintained borders (approximately at half of the borders), but there was a strong enrichment of cohesin at majority of maintained borders (Rad21, Nipped-B and Smc1) and Trl (Figures 4A, S4A and S8A). Furthermore, the maintained borders were enriched with Pol II, Mediator complex (MED30 and MED1) and Orc2. Significantly lower histone levels (H4/H3/H1) at maintained borders indicated the presence of highly accessible DNA (Figure 4B and Figure S8B). Noticeably, there is also strong divergent transcription at the maintained borders (Figure 4B). The lack of enrichment for Top2 at TAD borders that are maintained in the two mutants (Figure 4B) indicates a potential role for supercoiling at these borders.

**Figure 4.**
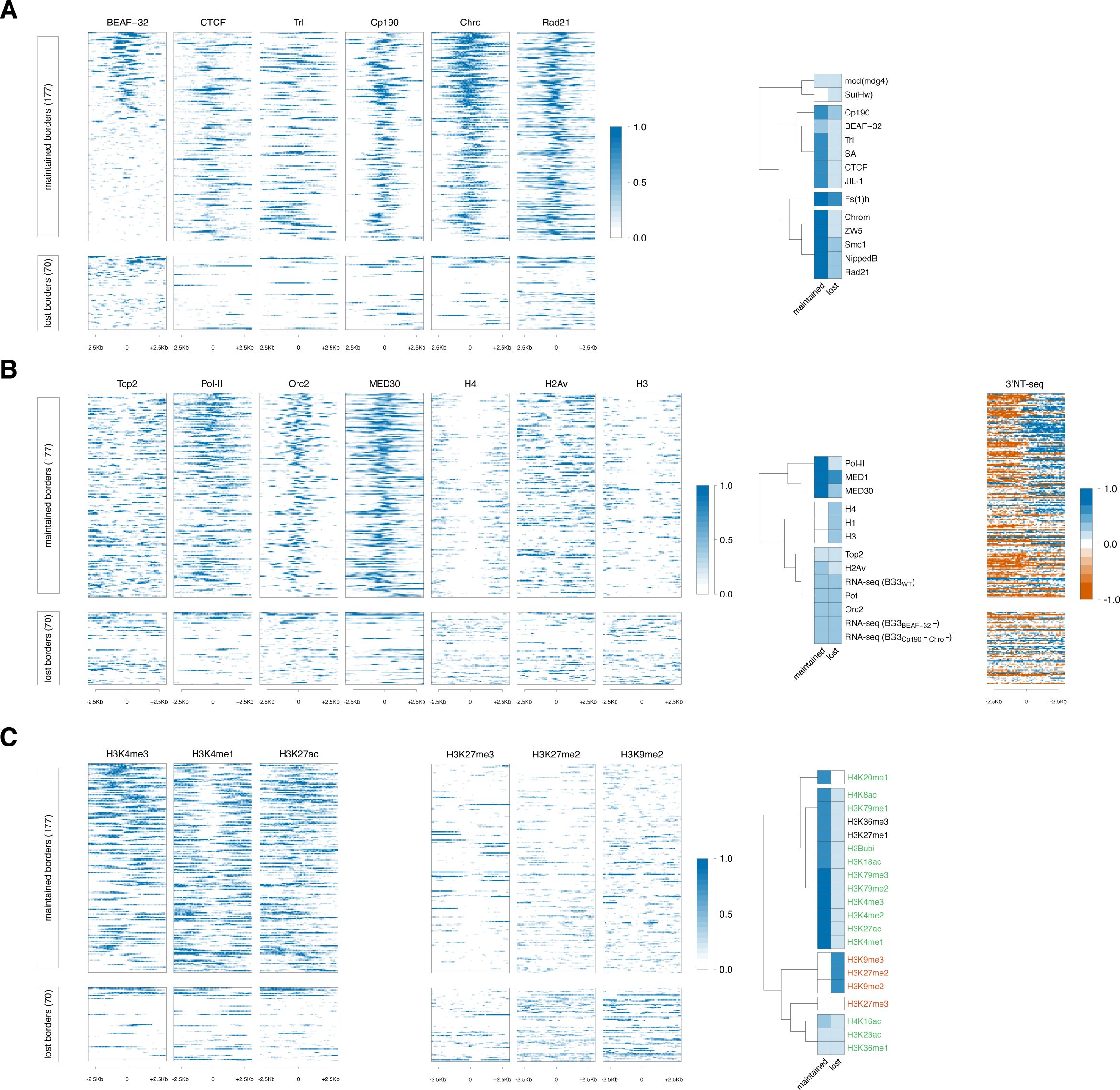
Chromatin features enrichment at TAD borders. (A) Profiles of architectural proteins (BEAF-32, CTCF, Trl, Cp190, Chro and Rad21) around direct maintained and lost TAD borders that were common in BEAF-32 knockdown and Cp190 Chro double knockdown. The right panel plots clustering of the signal at maintained and lost TAD borders (see Supplementary Materials and Methods). (B) Profiles of histones (H4, H3 and H2Av), transcription (Pol II, 3’NT-seq, MED30 and Top2) and replication (Orc2) at maintained and lost TAD borders. For nascent transcription, we used two colour schemes: orange for transcription on the negative strand and blue for transcription on the positive strand. (C) Profiles of histone modifications (H3K4me3, H3K4me1, H3K27ac, H3K27me3, H3K27me2 and H3K9me2) at maintained and lost TAD borders. We marked with red histone modifications associated with dense chromatin and with green histone modifications associated to open chromatin. Histone modifications that have been reported to be associated with both open and dense chromatin were marked by black. There is depletion of signal in the middle of the histone modifications heatmaps, which can be explained by the depletion of histones in those regions (see B).

The RNA-seq signal around maintained and lost TAD borders, does not show noticeable changes in the two mutants (Figure 4B and S8B). At maintained borders, given that there are negligible changes in gene expression, these results were expected (see Figure 3). Nevertheless, given the large number of differentially expressed genes associated to reorganised TADs (see Figure 3), one could expect a change in the RNA-seq signal at lost TAD borders in the mutant. Since the differentially expressed genes are randomly located inside the TAD (Figure S7), loss of TAD borders will often correlate with changes in gene expression at a larger distance from the TAD border and, this, cannot be captured in the analysis in the vicinity of TAD borders (Figures 4B and S8B).

By contrast, at lost TAD borders (in BG3_BEAF-32_^-^ and BG3_Cp190_^-^_Chro_^-^), there is less DNA accessibility and transcription indicating that these borders are in a repressed chromatin state (Figure 4B).

### Maintained borders are associated with active promoters and enhancers whereas lost borders are located in heterochromatin

Regulatory regions in the DNA can be defined by the presence of specific histone marks ^37^. Transcription has also been shown to be strongly implicated in the maintenance and formation of TADs ^6, 38, 39^. The presence of Pol II and nascent transcription at maintained borders and their absence from lost borders indicate the existence of two classes of TAD borders in *Drosophila*, active and repressed borders, which display different mechanisms of maintenance. A similar classification into active and repressed domains in *Drosophila* has been previously proposed^40, 41^. We investigated the presence of histone modifications to further dissect the potential factors and mechanisms that would be responsible for the maintenance of the TAD borders. We found that H3K4me3 (active promoter mark) and H3K4me1 (enhancer mark) together with H3K27ac were enriched at all maintained borders (Figure 4C). Interestingly, depletion of BEAF-32 from promoters and enhancers is not sufficient to result in the loss of these TAD borders, which indicates the presence of a redundant mechanism with a different protein(s).

We observed strong enrichment of MOF (involved in maintenance of H4K16ac), JHDM1 (H3K36me3 demethylase), ISWI and NURF301 (nucleosome sliding) and WDS (involved in maintenance of H3K4me3) preferentially at maintained borders (Figure S9). NURF301 was shown to co-localise together with Dref and Cp190^42^, which explains its enhanced level at the maintained TAD borders.

The lost borders were strongly enriched in H3K9me2, H3K9me3 and H3K27me2 (signatures for heterochromatin and Polycomb) suggesting a plausible association of these proteins with heterochromatin regions (Figures 4C and S8C). As we observed association of lost borders with heterochromatin and Polycomb, we further dissected and analysed the Polycomb complexes in detail at all borders. However, we did not observe enrichment of any Polycomb subcomplexes (Pc or dRING) at lost borders in the two mutants (Figure S9). Nevertheless, we did find enrichment of Su(var)3-9 and HP2, which explains the enrichment of heterochromatin at lost TAD borders in the two mutants (Figure S9). Note that, in Drosophila, Su(var)3-9 was previously reported to have a role in maintenance of TADs located in heterochromatin ^43^.

While we observed heterochromatic signatures at the lost borders (Figure 4C), previous research reported that TAD borders are mostly composed of euchromatin ^26, 38, 44–46^. Using a chromatin map in BG3 cells^47^, we investigated the chromatin states associated to maintained, new and lost TAD borders in each mutant. Our results confirm that indeed maintained, lost and new borders are enriched in enhancer and active TSS chromatin states (Figure S10A-C) and are depleted in heterochromatin (Figure S10A-C). In addition, lost borders also display partial enrichment in Polycomb state. This apparent difference in results at lost TAD borders can be explained by the fact that the analysis in Figure S10A-C is performed on TAD borders at a base pair resolution, while the analysis in Figures 4, S8 and S9 was performed over a 5 Kb region. When considering the same 5 Kb regions as in Figures 4, S8 and S9, one can observe an enrichment for heterochromatin and heterochromatin in euchromatin at lost TAD borders (Figure S10D). This means that while the majority of borders are enriched in enhancers or active TSSs, maintained borders are located in euchromatin and lost borders in heterochromatin.

One possibility is that lost borders, while euchromatic, display higher levels of Pol-II pausing. Using the Pol II pausing index definition from ^25^ (see Methods), we found only negligible differences in Pol II pausing for genes located within 5K windows around of maintained, lost and new borders (Figure S10E). This indicates that Pol II pausing does not differentially affect maintained or lost borders.

### A large proportion of maintained TAD borders in the knockdowns are also present in Kc167 cells and harbour housekeeping genes

Previously, we showed that Kc167 cells display more short-range interactions and fewer long-range contacts when compared to BG3 cells, which was true also after down-sampling to control for library size differences^26^. Given that the three mutants we analysed here display increased numbers of short-range contacts and reduced numbers of long-range contacts compared to WT BG3 cells (Figures 1A and 2A), this raises the question of how the 3D organisation of these mutants differs when compared to Kc167 cells. Our results show that there are significantly more short-range interactions and fewer long-range interactions in Kc167 cells compared to BG3_BEAF-32_^-^, BG3_Cp190_^-^_Chro_^-^ and BG3_BEAF-32_^-^_Dref_^-^ (Figure S11A). To further investigate the similarities between the BG3_BEAF-32_^-^, BG3_Cp190_^-^_Chro_^-^ and BG3_BEAF-32_^-^_Dref_^-^ and Kc167 cells, we compared the maintained, lost and new robust TAD borders in the mutants with the robust TAD borders in Kc167 cells. Approximately half of the maintained TAD borders in the three mutants are also strong TAD borders in Kc167 cells, but this decreases to less than 20% in lost and new borders (Figure S11B). This is true when comparing to both similar size ^26^ or significantly larger ^48^ Hi-C libraries in Kc167 cells. This indicates that nearly half of the maintained borders are housekeeping TAD borders, while the majority of lost borders are BG3 specific. The majority of genes present at the TAD borders conserved between Kc167_WT_, BG3_WT_, BG3_BEAF-32_^-^, BG3_Cp190_^-^_Chro_^-^ (176 out of 181) are housekeeping genes (Table S4 and Materials and Methods).

### Majority of chromatin loops in *Drosophila* are controlled by Mediator complex, Chro and Cp190

Chromatin loops represent enriched long-range 3D interactions and have been identified as important features in 3D chromatin organisation. In *Drosophila*, only a small number of loops have been detected^26, 48^. We identified loops in WT BG3 cells and in the three mutants and observed an increase in the number of loops in two mutants (BG3_BEAF-32_^-^ and BG3_Cp190_^-^_Chro_^-^) (Figure 5A). This could be attributed to the difference in sequencing depth between the different samples. Of the 770 loops that were detected in WT cells, in each mutant, approximately 200 are maintained and 300 maintain only one anchor in the same position (Figure 5B). We classified 140 loops that are maintained in both BG3_BEAF-32_^-^ and BG3_Cp190_^-^_Chro_^-^ mutants as maintained loops and 122 that are lost in both BG3_BEAF-32_^-^ and BG3_Cp190_^-^_Chro_^-^ mutants as lost loops (Figure 5C). Figure 5D confirms that the strong level of interactions is maintained in the two mutants at maintained chromatin loops, but this is not the case at lost loops. We also found that there is no statistically significant difference in the size of the lost and maintained chromatin loops (Figure 5E).

**Figure 5.**
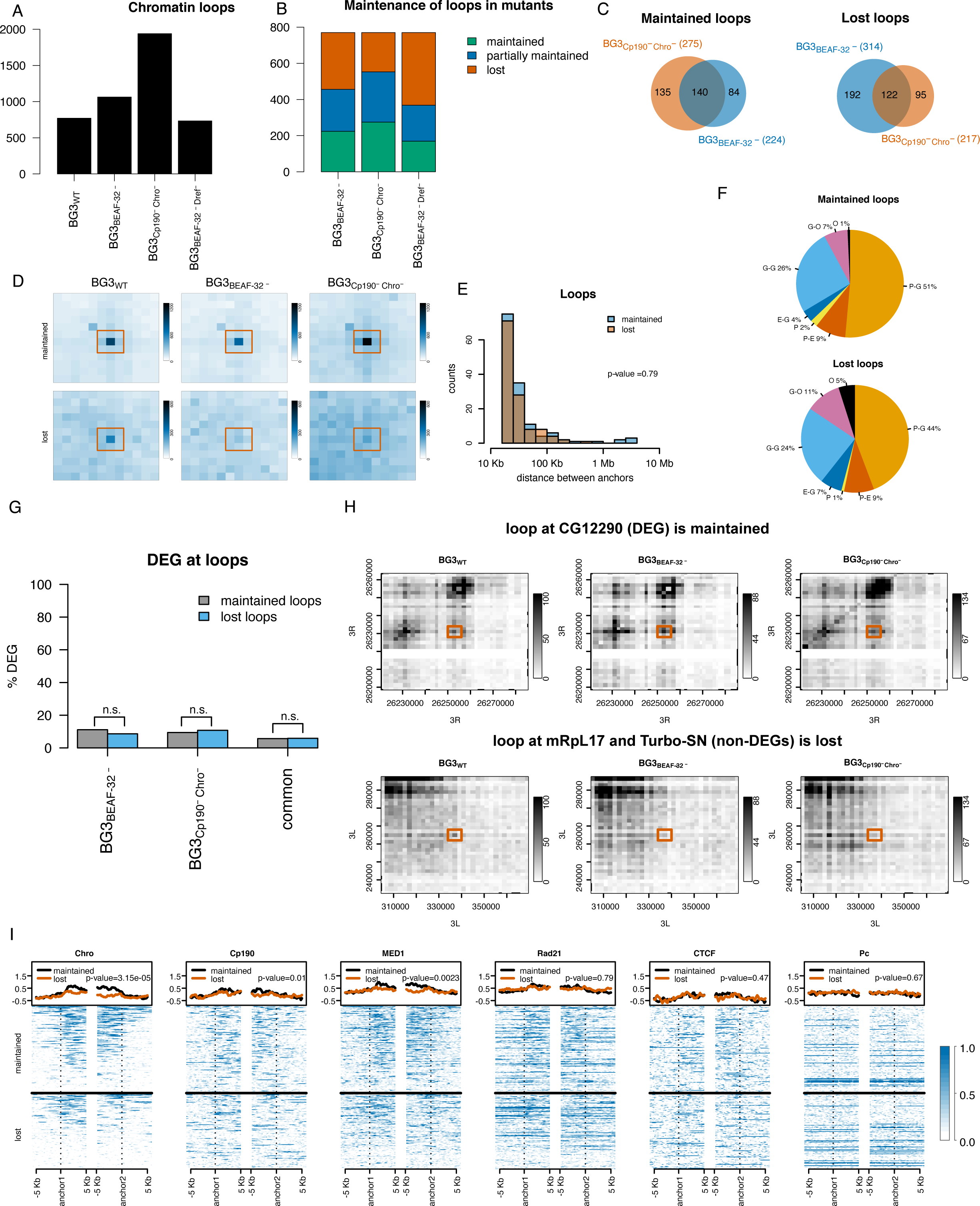
Chromatin loops. (A) Number of loops detected in WT and the three mutants. (B) Percentage of loops in the three mutants that maintain both of the anchors (maintained), only one of them (partially maintained) or lose both anchors (lost). (C) Overlap of loops maintained or lost between WT and BEAF-32 knockdown and between WT and Cp190 Chro double knockdown. We classify the commonly maintained borders in the two mutants as maintained and the commonly lost borders as lost. (D) Aggregate Peak Analysis (APA) using Juicer^72^ over the maintained (top) and lost (bottom) chromatin loops at 2 Kb resolution. (E) Size of the maintained and lost chromatin loops as defined in (C). We performed a Mann Whitney U test, which confirmed that the two distributions are not different. (F) Annotation of maintained and lost loops with respect to the features they connect: P - promoters (up to 1Kb upstream of TSS), E - enhancers, G - genes and O - others. We used STARR-seq for enhancer annotation^86^. (G) Percentage of genes that are differentially expressed and are associated with maintained and lost chromatin loops. We selected genes that have their promoter (up to 1Kb upstream of TSS) located at one of the anchors of the chromatin loops. There is no statistically significant difference between DEG at maintained and lost loops (Fisher’s exact test; p-value 0.37, 1.0 and 1.0). (H) Contact matrixes plots of a maintained (top) and a lost (bottom) loop. These maps were constructed with diffHic^87^ at 2 Kb resolution (the same used to detect loops) and contain 30 bins. Dark colour represents more contact. We scaled the pallet in the two mutants to account for to library size differences. (I) Enrichment of architectural proteins and transcription related factors at maintained and lost loops (Chro, Cp190, MED1, Rad21, CTCF and Pc). We performed a Mann-Whitney U test of the mean signal at maintained and lost loops (see corresponding p-values).

76% to 68% of these loops connect genes to each other or other genes (Figure 5F), indicating that they are involved in the formation of gene domains^28^. Only 9% of the maintained and lost loops are promoter-enhancer loops (Figure 5F), which indicates that this mechanism is less prevalent in *Drosophila* than previously proposed in mammalian systems^49^. When we select all genes that have their promoter located at one of the anchors of the loops, we found that only a small subset of genes (<10%) located at the lost or maintained loops display differential expression in the two mutants and this is true even when using a less stringent threshold to call differentially expressed genes (log_2_FC threshold of 1.0) (Figures 5G and S12). Furthermore, there is no statistically significant difference in the number of DEG at maintained and lost loops (Figures 5G). For example, a chromatin loop can be maintained in all three mutants while the target gene is differentially expressed (top panel in Figure 5H). Conversely, a lost chromatin loop can lead to no changes in gene expression of the target genes (bottom panel in Figure 5H). Thus, our results support a model where the presence or absence of a chromatin loop does not necessarily lead to regulation of the target gene.

Chro and Cp190 are known to be involved in long-range interactions^33^, but previous research identified the enrichment of Polycomb at *Drosophila* loops^48^. We found that both maintained and lost chromatin loops display high levels of BEAF-32 together with Chro and/or Cp190 at both anchors (Figure 5I and S13A); i.e., 92% of maintained and 84% of lost loops have binding of BEAF-32, Cp190 and/or Chro (Figure S13B). The maintained loops display higher levels of Chro at the anchors compared to lost loops, suggesting that the depletion of Chro (in BG3_Cp190_^-^_Chro_^-^) or blocking of its recruitment (in BG3_BEAF-32_^-^) is not sufficient to affect the maintained loops. In addition, 60% of lost loops lose binding of BEAF-32 and/or Cp190 upon their knockdown (Figure S13C-D), thus, providing support that these loops are lost as a direct consequence of the depletion of the architectural proteins in our mutants. Nevertheless, approximately half of the maintained loops lose BEAF-32 and/or Cp190 upon knockdown (Figure S13C-D), which suggests that Chro is recruited by additional proteins at maintained loops or that other factors could help maintain these loops (Figure S13A). We observed an enrichment of MED1 at the anchors of maintained and lost loops, but also enrichment of CTCF and cohesin subunit Rad21. The majority of chromatin loops in our dataset are located near a MED1 ChIP peak (Figure S13A), indicating that Mediator complex would be more important for chromatin loops in *Drosophila*. We also observed a small number of loops with enrichment of Polycomb peaks near their anchors (Figure 5 and S13), but this is less pronounced than in the case of Mediator complex.

### BEAF-32, Cp190, Chro and Dref knockdown does not affect A/B compartments

The checkerboard pattern seen on Hi-C maps led to the identification of A and B compartments which mark active and inactive regions of chromatin ^50^. A/B compartments were also identified in *Drosophila*^39^ using 10 Kb bins, and we showed that compartmentalisation changes between cell lines^26^. The current working model assumes that compartments harbour several TADs and they display different mechanisms for maintenance compared to TADs. To investigate if the changes in the TADs lead to changes in the A/B compartmentalisation of the genome, we computed the A/B compartments at 10 Kb resolution in the three mutants (see Methods). Our results confirm that there are negligible changes in the proportion of the genome that is in the A or B compartments in all mutants (Figure S14A). Nevertheless, we identified some switching between the A and B compartments (less than 5%) (Figure S14B). When we zoomed in, we observed that the majority of these compartments are robust and consistent in the WT and mutants (Figure S14C). One interesting observation is that there is some rare local spreading of the B compartment (heterochromatin) into the A compartment (euchromatin) (e.g., yellow stripe in Figure S14C).

Saddle plots confirm that regions belonging to the same compartments (lower right corner A-A interactions and upper left corner B-B interactions) are enriched in interactions, while regions belonging to different compartments (lower left corner A-B interactions and upper right corner B-A interactions) are depleted in interactions (Figure S14D). Compartments strengths are similar, with only a small decrease for BEAF-32 single knockdown. This confirms that BEAF-32, Cp190, Chro and Dref have little effect on compartmentalisation. Altogether, our results indicate that organisation of compartments in Drosophila is independent of the organisation of TADs.

When investigating in which compartments are TAD borders localised, we found that most of the maintained borders are localised in the A compartment, while most of the lost or new borders are localised in the B compartment (Figure S15A). This is not surprising since most of the lost borders are located in repressed chromatin, while the maintained ones are in active chromatin.

The majority of compartments that switch do not harbour any DEGs, even when using a lower threshold to call differential gene expression (log_2_FC threshold of 1) (Figure S15B). Furthermore, the fact that a compartment contains DEGs does not mean that all genes in that compartment change expression in the same direction (either upregulated or downregulated). For example, spreading of B compartment in Figure S14C corresponds to three genes displaying different behaviours: *ine* gene is downregulated, *Dp* is upregulated and *FIG4* maintains expression in all three mutants (all three genes are located within the yellow stripe in Figure S14C). The relationship between changes in gene expression and compartment switching is complex and often compartment switching cannot be explained by a majority of genes changing expression in the same direction. Note that RNA-seq libraries capture only polyA transcripts and do not include other transcripts such as eRNAs or lncRNAs which could potentially contribute to compartment switching.

## Discussion

The enrichment of architectural proteins at TAD borders raises the question of whether they have a functional role in TAD organisation or whether their co-localisation with borders is correlative in nature. In mammalian systems, depletion of CTCF or Cohesin disrupts TADs^21–23^. In *Drosophila*, several architecture proteins (including BEAF-32, Chro and Cp190) are enriched at TAD borders, but their functional role at TAD borders has not previously been investigated^16, 24–27, 45^. Our results confirm that the architectural proteins are essential for TAD borders and their depletion results in reorganisation of TADs. In particular, we found that TAD borders mainly found in heterochromatin are lost upon depletion of BEAF-32 or Cp190 and Chro. Cp190 and Chro cannot bind independently to DNA, but are recruited, mainly, by BEAF-32^33^. The majority of the lost borders are common between the BEAF-32 mutant and Cp190 and Chro double mutant, but there are also borders that are specific to each mutant. Furthermore, we identify a subset of TAD borders that are not affected by the depletion of BEAF-32, Cp190 and Chro. These borders are enriched in cohesin and Mediator complex and also in CTCF and Trithorax-group (Fs(1)h, NURF301, ISWI, mod(mdg4), ASH-1 and Trl). This supports a model where several complexes are redundant and can compensate for the loss of BEAF-32, Cp190 or Chro. However, 70% of TAD borders that are maintained have retained binding of BEAF-32 and/or Cp190 upon the depletion of these architectural proteins (see Figure S4B-C).

Finally, Dref shares a similar binding motif to BEAF-32, which suggests that it could potentially replace it following BEAF-32 knockdown. Our BEAF-32 Dref double mutant results in a larger number of TAD borders being lost, supporting the model in which Dref compensates the loss of BEAF-32. The borders that are specifically lost in BEAF-32 and Dref double knockdown are borders displaying binding of BEAF-32, Cp190 and/or Chro and, thus, Dref would provide redundancy for partial loss of BEAF-32.

To investigate that the effects we observe in 3D chromatin organisation are not a reflection of cell cycle arrest ^46, 51^, but are due to the knockdown of architectural proteins, we have performed a FACS analysis. This showed that none of our knockdowns lead to changes in the cell cycle progression (Figure S1E), thus, confirming that the changes in 3D chromatin organisation are not caused by cell cycle arrest.

Altogether, our results confirm that, while the majority of TAD borders are enriched in enhancers or active TSSs, there are two classes of TAD borders: *(i)* TAD borders located in euchromatin and *(ii)* TAD borders located in heterochromatin. While the former are maintained upon depletion of BEAF-32, Cp190 and Chro, the latter are lost. This classification of TAD borders is additionally supported by the preferential localisation of maintained borders in A compartment (active chromatin) and of lost and new borders in B compartment (repressive chromatin) (see Figure S15A).

The enrichment of divergent transcription at BEAF-32 enriched TAD borders that are maintained in the two mutants, when coupled with the lack of enrichment for Top2 at these borders, possibly indicates that negative supercoiling accumulates at these TAD borders, which may be due to active transcription. This negative supercoiling is not relaxed due to lack of Top2. When negative supercoiling accumulates at these borders, positive supercoiling may accumulate inside TADs, which indicates a role for supercoiling in TAD borders^52, 53^.

### TAD reorganisation and transcription

We identified between approximately 600 and 800 differentially expressed genes in the three mutants and the majority of those are located within TADs that lose one or both borders or shifted the position of the borders (more than 89%). We also found that there are more statistically significant DEGs than expected by chance in reorganised TADs, however this is mainly the case when TAD borders move more than 2 Kb away from their WT position. This indicates that usually strong TAD reorganisation is coupled with significant changes in gene expression. Nevertheless, there are also examples where discrete changes in TAD borders correspond to changes in gene expression ^54, 55^. For Cp190 and Chro double knockdown, we did not see a statistically significant association between DEGs in reorganised TADs. This can be explained by the fact that these two proteins are also recruited to the DNA by other proteins that would not be involved in TAD border organisation^56^. In this case, a subset of DEGs in Cp190 and Chro double knockdown are not associated to reorganisation of TADs and, thus, the statistical significance of association of DEG and reorganisation of TADs is reduced.

We also observed more upregulated genes than downregulated, which suggests that TADs have a role in maintaining a repressed state of chromatin. Downregulation of genes in these mutants can be explained by the loss of TAD borders in heterochromatin. Previous work in *Drosophila* did not identify any connection between changes in TADs and changes in gene expression^34^. These contradicting results can be explained by the stronger reorganisation of the TADs in our mutants compared to the TADs reorganisation observed on the balancer chromosomes. Recently, it was shown that there are significant changes in gene expression corresponding to reorganisation of TADs in human cancers, but only 14% of changes in TAD organisation result in strong changes in gene expression (more than twofold)^13^. Our findings are consistent with these results and emphasise that the functional role of TAD organisation is conserved between species.

One question that is still unanswered is whether the changes in gene expression are caused by the changes in TAD organisation or whether depletion of architectural proteins affects transcription, causing the observed changes in TAD organisation. Previous studies showed that TADs appear together with transcription activation in the Drosophila zygote indicating a functional role of transcription in TAD formation, but blocking transcription elongation only marginally affects TADs in the Drosophila embryo^45^. Furthermore, a 10-20 fold activation of transcription using the CRISPR/Cas9 system in mouse neuronal progenitor cells was not sufficient to induce TAD boundary formation^57^. Results from these alternative approaches suggest that changes in gene expression do not lead to reorganisation of TADs, but further work is needed to confirm this.

### Chromatin loops and gene regulation

Our analysis revealed that the chromatin loops in *Drosophila* can be classified in three large classes: *(i)* BEAF-32 with Chro and/or Cp190, *(ii)* Mediator complex and *(iii)* Polycomb (Figures 5D and S13). Previous work reported that chromatin loops in *Drosophila* are controlled by Polycomb^48^, but our results show that Polycomb loops are just a small subset compared to Chro/Cp190 and Mediator complex loops. Depleting Chro/Cp190 or BEAF-32 (protein that recruits Chro/Cp190 to DNA) results in the loss of loops, mainly those loops displaying weaker Chro/Cp190 enrichment, suggesting concentration dependent control. Chro and Cp190 were shown to be involved in long range interactions in *Drosophila* ^33^ and our results confirm that the majority of chromatin loops in *Drosophila* are controlled by these proteins. We also found enrichment of cohesin and CTCF at chromatin loops, indicating that they might have a role in chromatin loop formation in *Drosophila*^18, 58^. In particular, half of maintained loops lose BEAF-32 and/or Cp190 binding upon knockdown, indicating that CTCF and cohesin could play a role in the maintenance of these loops.

Some interactions between specific DNA regions identified in the contact maps are shown to arise from promoter-enhancer loops (a chromatin loop having an enhancer at one end and a promoter at the other)^49^. In *Drosophila* BG3 cells, we found that only 10% of the chromatin loops are promoter-enhancer loops and one possible explanation for this is that the annotation of enhancers is not comprehensive^59^. Even if this is the case, only approximately half of the loops have promoters at one end indicating that majority of interactions are not regulatory in nature ^60–62^. Furthermore, even in the case when a promoter has a 3D contact with a regulatory sequence, less than 10% of genes display differential expression when the contact is lost, but the same is true at maintained loops. This suggests that the presence of chromatin loops would not be essential for controlling gene transcription in the majority of cases ^58, 63–65^.

## Materials and Methods

### Cell Culture and Knock down

*Drosophila* BG3 cells were cultured at 25°C in Schneider’s insect medium (Sigma), supplemented with 10% FBS (Labtech), 10 mg/l insulin (Sigma, I9278) and Antibiotic Penstrep. Primer sequences for Cp190, Chro and BEAF-32 dsRNAi were obtained from the *Drosophila* RNAi Screening Center database (http://flyrnai.org) (see Table S5). The primers with T7 promoter sequence were used to amplify the IVT templates from wild type genomic DNA using Dream Taq DNA Polymerase Kit (Thermo Scientific, EP0703). The PCR products were checked by electrophoresis and purified using a Fastgene PCR Purification Kit (Fastgene). The purified PCR products were then used as templates to synthesis dsRNA using the MEGAscript T7 Kit (Life Technologies, AM1334) according to the manufacturer’s recommendations. The BG3 cells were transfected with 50ug of dsRNA using Fugene (Promega) according to manufacturer’s protocol. Cells were harvested after 72 hours and processed for downstream experiments accordingly.

### Western blot

Cells were pelleted, washed in PBS and resuspended in SDS PAGE loading buffer at a concentration of 40 000 cells per µl, sonicated and boiled for 4 minutes. 5µl of lysate were loaded on a 10% polyacrylamide gel. The proteins were transferred on nitrocellulose and saturated 1 hour with 5% skimmed milk (or 1% BSA for the Chro antibody) in PBS tween 0.1%. the blots were incubated overnight with anti-BEAF-32^66^ (1/200) anti-Chro^67^ (1/200) anti-Cp190^68^ (1/5000) or anti-Dref^32^ (1/5000). Secondary antibody (peroxidase anti rabbit for Cp190 and Dref and peroxidase anti mouse for BEAF and Chro) were incubated at a 1/10000 dilution. They were visualised with Pierce ECL western blotting substrate using the Fujifilm LAS4000 gel imaging system. anti-BEAF-32 and anti-Chro (12H9-4A2) were purchased from Developmental Studies Hybridoma Bank, while anti-Cp190 and anti-Dref were kindly provided by Dr Rob White and Dr Professor Masa Yamaguchi respectively.

### FACS

Cells were pelleted, washed in PBS and resuspended in 50% ethanol in PBS and stored until analysis at 4^0^C. On the day of the analysis cells were pelleted, washed in PBS and resuspended in FACS PI buffer (PBS, 01% triton, 100µg/ml RNase and 50µg/ml propidium iodide) at a concentration of 10^6^ cells/ml. The cell cycle profile was analysed with the Guava easycyte HT flow cytometer using the Incyte software and FlowJo. For each sample 15000 cells were analysed.

### In situ Hi-C protocol

Hi-C libraries were generated from 10 million cells by following the *in situ* Hi-C protocol as mentioned in^26^. Briefly, crosslinked cells were lysed and genome was digested using DpnII (NEB) overnight. The overhangs were filled with Biotin-16-dATP (Jena Bioscience) followed by ligation and de-crosslinking with proteinase K digestion. The sample was further sonicated using Bioruptor. Biotinylated DNA was pulled down using Dynabeads MyOne Streptavidin T1 beads (Life technologies, 65602). Selected biotinylated DNA fragments ranging from 200-500bp were then ligated with illumina adaptors (NEB). The libraries obtained from biological replicates were multiplexed and further sequenced at Oxford Genomics Centre and Edinburgh Genomics (Genepool) using HiSeq4000.

### Hi-C analysis

Each pair of the PE reads was aligned separately to *Drosophila melanogaster* (dm6) genome^69, 70^ using BWA-mem^71^ (with options -t 20 -A1 -B4 -E50 -L0). HiCExplorer was used to build and correct the contact matrices and detect TADs and enriched contacts^25^. The contact matrices were built using the DpnII restriction sites. We also used 100 Kb bins for plotting Figures S3 and S8 only and 10Kb for compartments^39^. Using a minimum allowed distance between restriction sites of 150 bp and a maximum distance of 1000 bp, we obtained a matrix with 217,638 bins with a median width of 529 bp. After filtering, we obtained between 18 M and 65 M valid pairs (see Table S1). Note that the number of reads and valid pairs used in this study are within values successfully used for previous work in Drosophila cells to detect TADs, chromatin loops and compartments (e.g., ^14, 26, 46^). In addition, we also showed that these libraries are robust to downsampling (Figure S3)^26^. The matrices were corrected using the thresholds in Table S2, where values were selected from the diagnostic plots (Figure S16). By using the corrected contact matrices, we detected TADs of at least 5 Kb width using a P-value threshold of 0.01, a minimum threshold of the difference between the TAD-separation score of 0.04, and FDR correction for multiple testing (--step 2000, --minBoundaryDistance 5000 --pvalue 0.01 --delta 0.04 -- correctForMultipleTesting fdr). We selected these parameters to ensure that we recover a similar number of TADs as previously reported^26^. Finally, we called strong TAD borders using a stringent value of the threshold of the difference between the TAD separation score of 0.08. This value ensured that we retrieved the strongest half of TADs. The enriched contacts were extracted with HiCExplorer using the observed/expected ratio method.

### Chromatin loops

Chromatin loops were called with the HICCUPS tool from the Juicer software suite^72^ on all mutants as done previously^26^. Loops were called using a 2 kb resolution, 0.05 FDR, Knight-Ruiz normalisation, a window of 10, peak width of 5, thresholds for merging loops of 0.02,1.5,1.75,2 and distance to merge peaks of 20 kb (-k KR -r 2000 -f 0.05 -p 5 -i 10 –t 0.02,1.5,1.75,2 -d 20000).

### Compartments

Compartments were called as described in^26, 28, 50^. More specifically, we used Juicer^72^ to compute the eigenvectors in 10 Kb bins for all conditions^26^. The sign of the correlation between the GC content and eigenvectors was used to flip the sign of the eigenvector^73^. Bins with negative eigenvalues were assigned as a B compartment, while bins with positive eigenvalues were assigned as an A compartment. Chromosomes 4 and Y are relatively small making the compartments call difficult and, thus, we excluded them from the compartment analysis.

### Saddle Plot and Compartmentalisation Strength

We use the procedure similar to ^74^. We rank each genomic region by their eigenvector value over 30 percentile bins. Note that we only included regions that fall in the [2.5%, 97.5%] quantile interval to eliminate the effect of outliers. We then calculate the mean value over intra-arm Pearson correlation values between regions with different percentiles. To make matrices comparable and generate saddle plots, we normalised averaged matrices using the absolute maximum values over WT and all three mutants. The compartment strength is calculated as the ratio of homotypic A-A and B-B interactions to heterotypic A-B and B-A interactions ^75^. The ratio is calculated using the averaged signals over corner sub-matrices of 10×10 bins. Note, that the compartment strength ratio uses non-normalised signal.

### Definition of housekeeping genes

We identified 113 strong TAD borders that are conserved between Kc167^26^, BG3_WT_, BG3_BEAF-32_^-^, BG3_Cp190_^-^_Chro_^-^ and 186 genes that are within 5 Kb of these borders. We then identified expression levels for 181 of them in 85 samples (tissues, cells, conditions or developmental stages)^76^ and classified genes as housekeeping if their expression was in the top 40^th^ percentile in all 85 samples^77^.

### RNA extraction and sequencing

RNA extraction was carried out using Trizol according to manufacturer’s instructions. RNA was further DNase treated and purified using RNeasy Mini kit (Qiagen) following the manufacturer’s protocol. RNA was assessed qualitatively and quantitatively using Quibit and Bioanalyzer 2100(Agilent). PolyA RNA selection, library preparation and sequencing were carried out by Novogene.

### RNA-seq analysis

Reads were first trimmed using Trimmomatic (v0.39)^78^ and then aligned to the *Drosophila melanogaster* (dm6) genome ^69, 70^ using TopHat(v2.1.2)^79^ with Bowtie2(v2.3.4.1)^80^ (Table S3). Finally, we used Picard tools (http://broadinstitute.github.io/picard/) to deduplicate reads, HTseq^81^ to count reads and then DESeq2^82^ to detect differential expressed genes. For DESeq2 we selected transcripts with at least 10 reads and used a p-value threshold of 0.05 and a log_2_FC threshold of 2.0 (for compartments and loops we reduced the log_2_FC threshold to 1.0). A previous work used Affymetrix GeneChip expression analysis to quantify changes in transcription upon BEAF-32 knockdown in BG3 cells and they observed negligible changes in gene expression^56^. Using RNA-seq, we found a larger number of genes displaying differential expression, but this can be explained by the increase sensitivity of RNA-seq.

### Analysis of Differentially and Non-Differentially Expressed Genes

We removed all genes that were not expressed in WT or any of the mutants and then we split the genome on short regions belonging to single WT TADs or mutant TADs. Each region was classified as follows: *(i) conserved 2 borders* if both borders of WT TAD that contains this region are conserved, *(ii) conserved 1 border* if only one of the borders moved more than 2 Kb compared to WT position, *(iii) mutant specific borders* if both borders moved more than 2Kb compared to their WT position and *(iv) fuzzy borders* if both borders moved less than 2Kb compared to their WT position. We then performed a permutation test using regioneR package with 1000 permutations (Gel et al., 2016).

### Pol-II pausing index

We followed the method from ^46^ and computed the pausing index as the ratio of the mean Pol-II ChIP signal over the promoter and over the gene body. Promoter region was selected from 200 bp downstream to 50 bp upstream of TSS and gene body from 50 bp upstream to gene end. Values of 0 and below were discarded.

### Data

The full list of datasets used can be found in Supplementary Tables S6-S11.

*ChIP-chip:* We used the ChIP-chip datasets generated and pre-processed (M values smoothed over 500 bp) by the modENCODE Consortium. The Fs(1)h, MED1, MED30, NippedB, Rad21, SA and Smc1 ChIP-chip datasets were downloaded from^83^. To call peaks for MED1 and Rad21, we first trimmed the reads using Trimmomatic^78^ (0.38), merged the two replicates (38.4M and 16.7M reads respectively), aligned them to the genome using bowtie2^80^ (using default parameters and achieving >94% alignment rate) and then used macs2^84^ for peak calling (q-value of 0.05 and using the corresponding input ChIP).

In some cases, we merged several ChIP peaks datasets: BEAF-32 (GSE32775, GSE20811, GSE32773 and GSE32774), Cp190 (GSE32776, GSE20814 and GSE32816) and CTCF (GSE20767, GSE32783 and GSE32782).

*DNase-seq:* We used pre-processed DNase-seq profiles from the modENCODE Consortium^37^.

*3’NT-seq:* We used pre-processed 3’NT-seq in BG3 cells (GSE100545) from^85^.

## Supplementary materials and methods

Information on the comparison of borders between WT and mutants, analysis of chromatin signals at TAD borders and clustering analysis can be found in *Supplementary materials and methods*.

## Data access

All Hi-C and RNA-seq datasets from this study have been submitted to the NCBI Gene Expression Omnibus (GEO; http://www.ncbi.nlm.nih.gov/geo/) under accession number GSEXXXXXX. The pipeline for Hi-C data analysis and RNA-seq is available as Supplemental Code.

## Acknowledgements

We would like to thank Dr Rob White and Dr Pradeepa Madapura for useful discussion and comments on the project and the manuscript. We also want to thank Zabet lab (especially Olivia Grant, Romana Pop and Bhavana Kayyar) for useful comments and discussions on the project and the manuscript. We would like to thank Professor Masa Yamaguchi and Dr Rob White for kindly sharing anti-Dref and anti-Cp190 respectively. The analysis was performed on the HPC at University of Essex and we would like to thank Stuart Newman for his support on using the cluster.

## Author Contributions

K.T.C. and N.R.Z. conceived and designed the experiments. K.T.C., G.C., I.H. and S.B.D performed the experiments. L.A.M., J.C.W. and N.R.Z. analysed the data. N.R.Z. and S.C. supervised the experiments. N.R.Z., H.D. and A.H. supervised the data analysis. K.T.C., L.A.M., G.C., S.C., H.D., A.H. and N.R.Z. wrote the paper.

---

**Figure S1.**
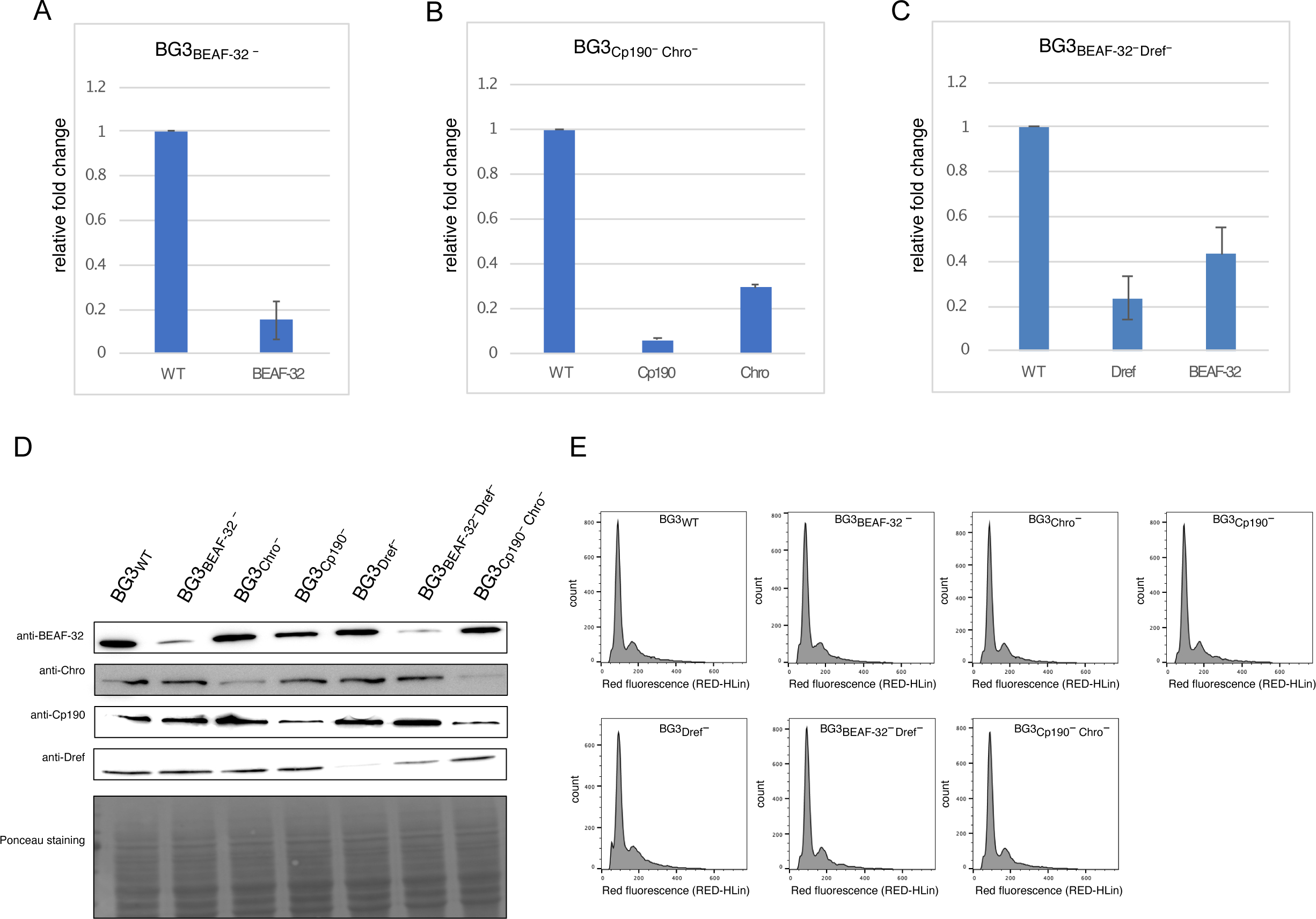
RNAi efficiency and cell cycle analysis in the three mutants. RT-qPCR in (A) BEAF-32 single knockdown, (B) Cp190 Chro double knockdown and (C). BEAF-32 Dref double knockdown. Data is normalised with Rpl32 and plotted relative to WT levels. (D) Western Blots showing the depletion of BEAF-32, Chro, Cp190 and Dref following RNAi treatment in WT and BEAF-32, Cp190, Chro and Dref single and double knockdowns as indicated in the panel. (E) FACS analysis showing no effect on cell growth or cell cycle arrest in WT or BEAF-32, Cp190, Chro and Dref single and double knockdowns.

**Figure S2.**
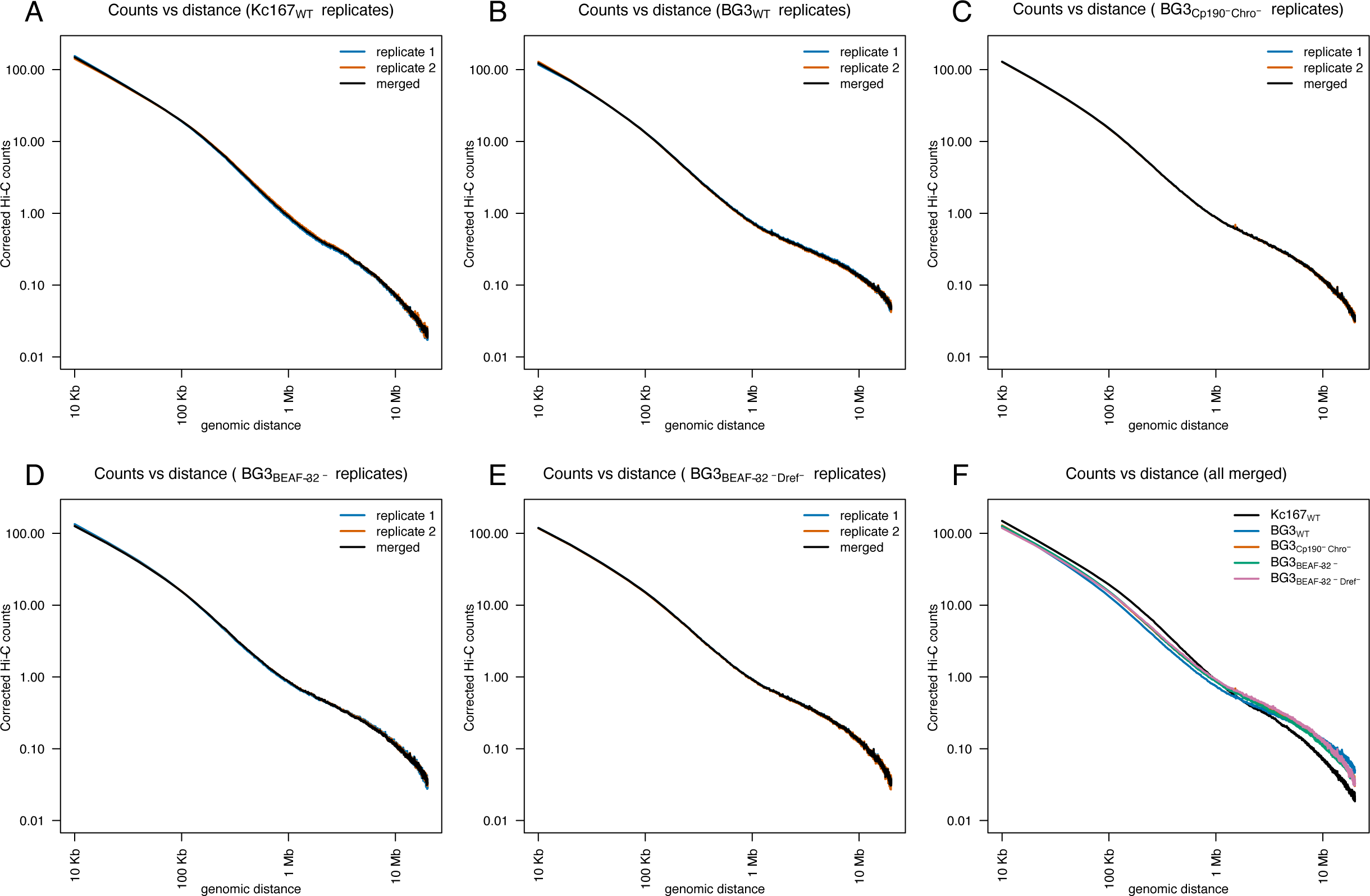
Counts vs distance plots for Hi-C datasets. For each Hi-C dataset we generated two replicates. In plots (A-E) we compare the two replicates with a merged dataset of the two replicates for (A) Kc167 WT cells, (B) BG3 WT cells, (C) BG3 cells with Cp190 Chro double knockdown, (D) BG3 cells with BEAF-32 single knockdown and (E) BG3 cells with BEAF-32 Dref double knockdown. The plots confirm that there are negligible differences between the two replicates and the merged dataset in each condition. In (F) we plot the merge datasets for all five conditions. The plot shows that the three mutants in BG3 display different behaviours to WT, but not as drastic as the differences between the BG3 and Kc167 cells.

**Figure S3.**
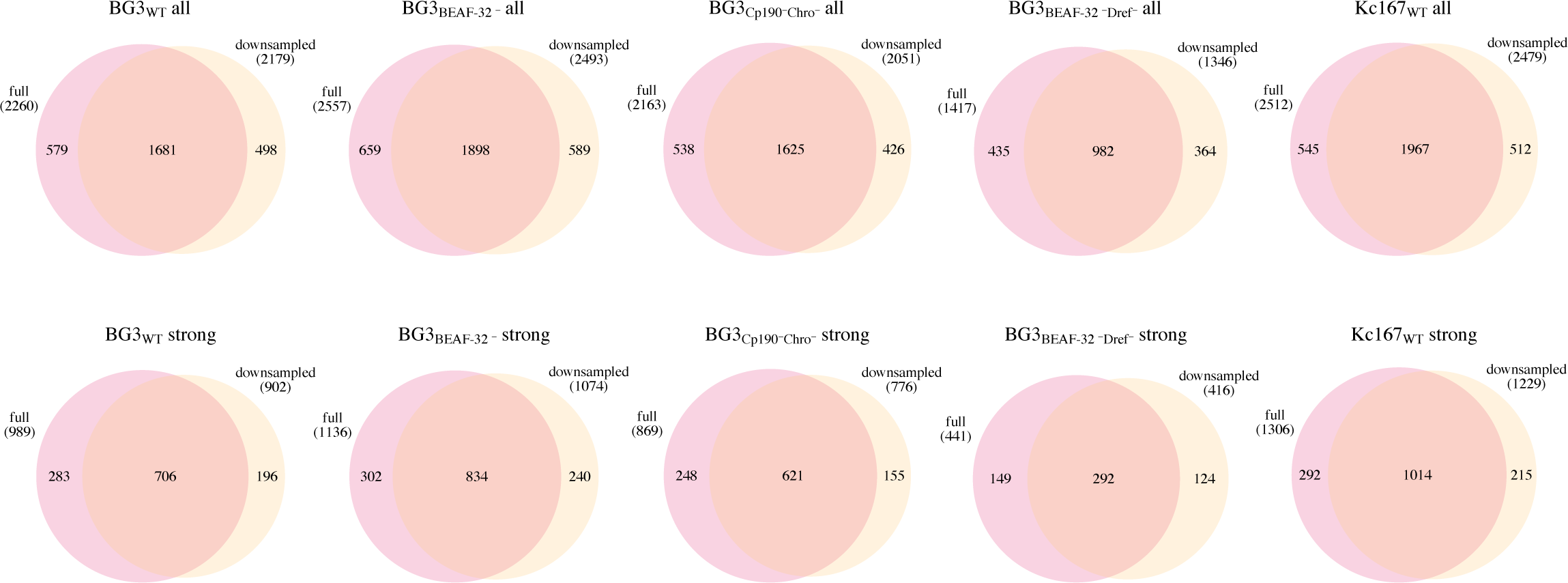
Robustness of TAD borders. We considered both the full dataset and a downsampled dataset where we randomly removed 20% of the reads. We consider the case of all (top) and strong (bottom) borders separately for Hi-C datasets in BG3 WT and the three mutants: BEAF-32 single knockdown, Chro and Cp190 double knockdown and BEAF-32 and Dref double knockdown. We also consider the case of Kc167 WT cells.

**Figure S4.**
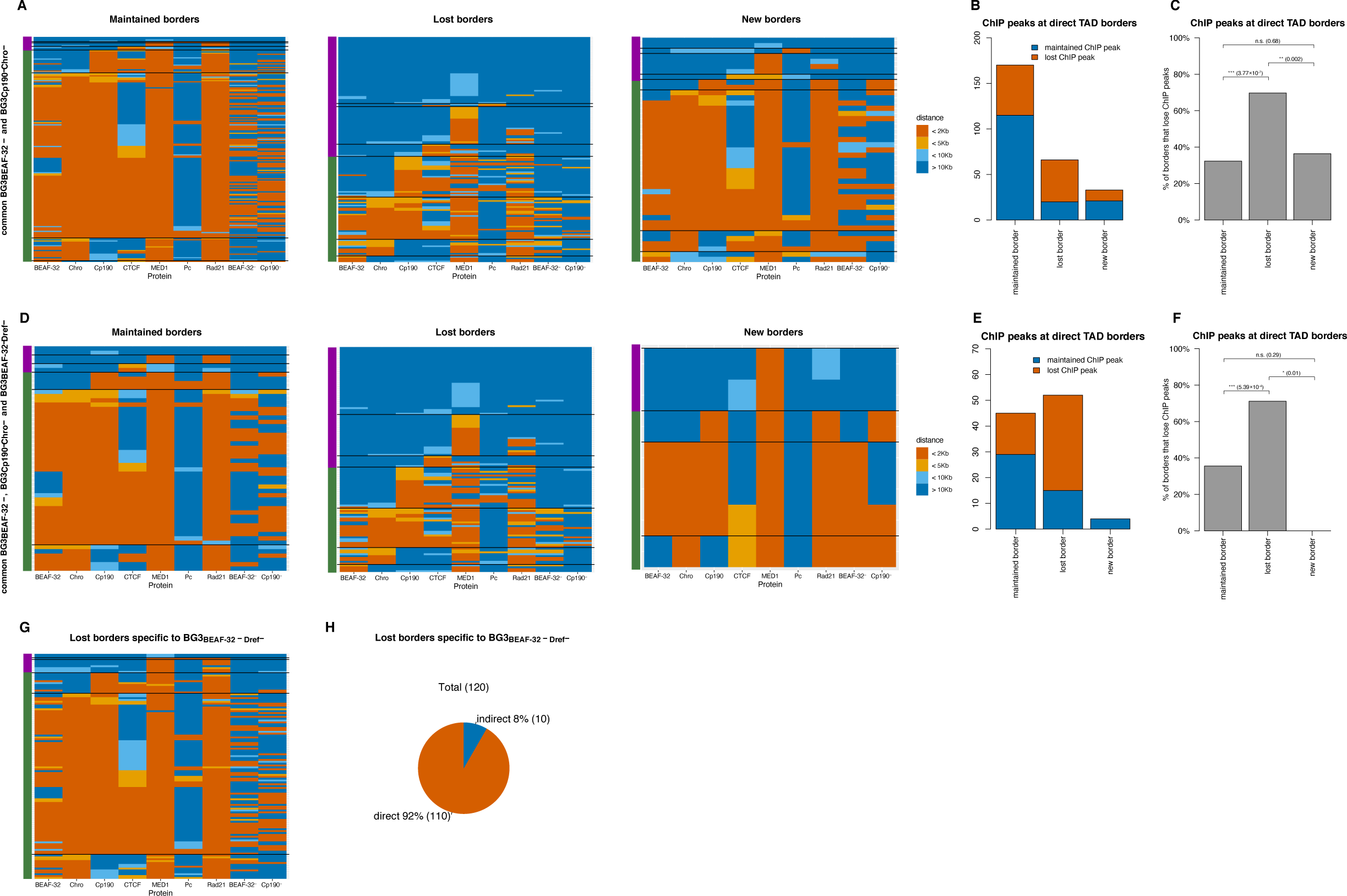
Direct and indirect TAD borders. We considered separately the cases of maintained, lost and new borders that are: (A-C) common between BEAF-32 single knockdown and Cp190 and Chro double knockdown and (D-F) common between BEAF-32 single knockdown, Cp190 and Chro double knockdown and BEAF-32 and Dref double knockdown. (A and D) Heatmaps plotting the distance of the closest ChIP peak from a maintained, lost and new border for: BEAF-32 (WT and BEAF-32 knockdown), Chro (WT), Cp190 (WT and Cp190 knockdown), CTCF (WT), MED1 (WT), Pc (WT) and Rad21 (WT). Green bar on the side of each heatmap marks the direct borders (borders that show binding of BEAF-32, Chro and/or Cp190), while purple indirect borders (all other borders). (B and E) number of TAD borders that have BEAF-32 or Cp190 ChIP in WT cells and lose or maintain those peaks in BEAF-32 and Cp190 single knockdowns. (C and F) percentage of TAD borders that have BEAF-32 or Cp190 ChIP in WT and lose them in the in BEAF-32 and Cp190 single knockdowns. We performed a Fisher’s exact test and the corresponding p-value is displayed above the barplots. (G) Heatmap plotting the distance of the closest ChIP peak from lost TAD borders that are specific to BEAF-32 and Dref double knockdown for: same proteins as in (A and D). Green bar on the side of each heatmap marks the direct borders (borders that show binding of BEAF-32, Chro and/or Cp190), while purple indirect borders (all other borders). (H) Total number and proportions of lost TAD borders that are specific to BEAF-32 and Dref double knockdown and have binding of BEAF-32, Cp190 and/or Chro.

**Figure S5.**
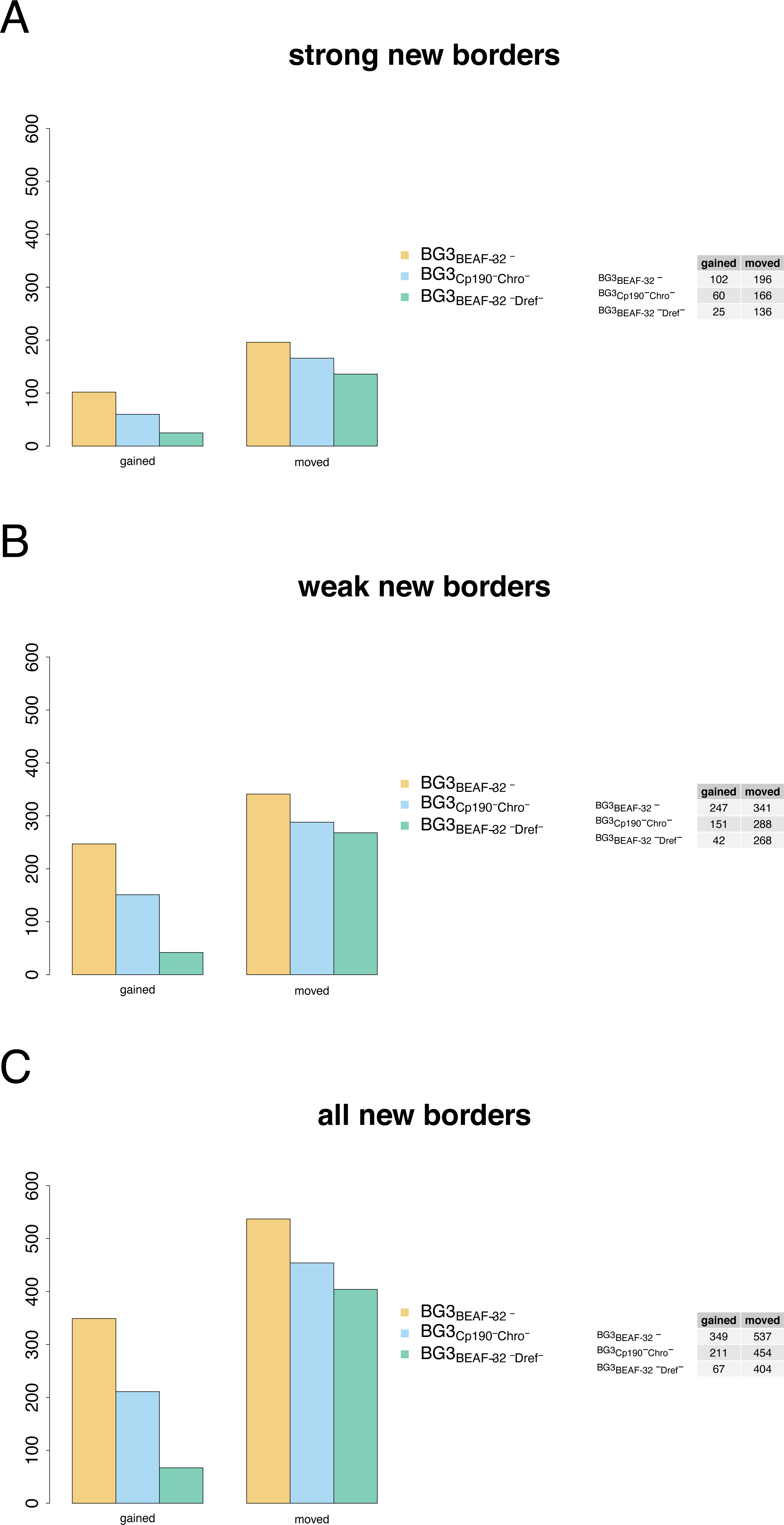
Classification of new TAD borders in the mutants. New borders in the three mutants (BEAF-32 knockdown, Cp190 Chro double knockdown and BEAF-32 Dref double knockdown) can be gained as a consequence appearing inside a WT TAD (splitting a TAD) or moved when they correspond to relocation of a WT TAD border. We considered the cases of: (A) new strong borders; (A) new weak borders; (C) all new borders.

**Figure S6.**
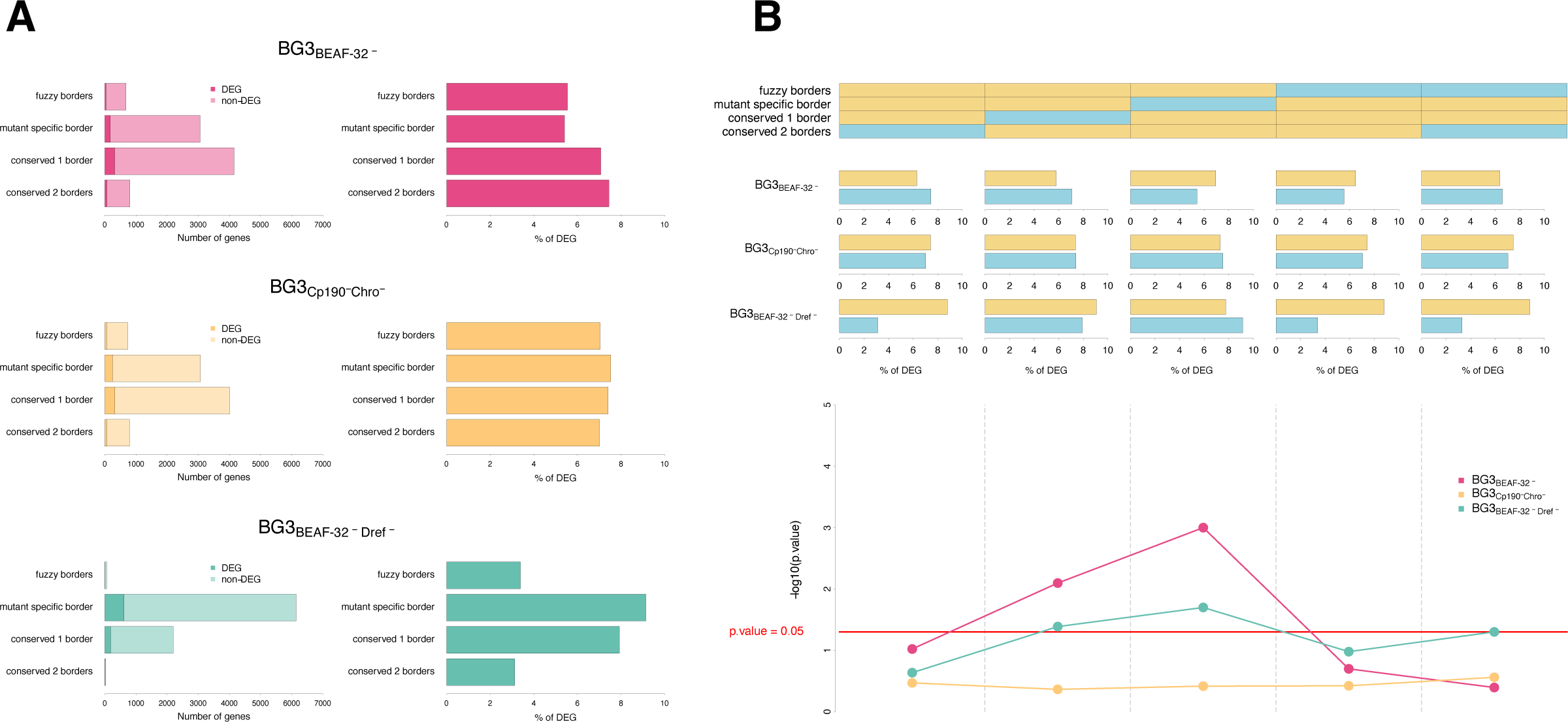
Comparison of differentially and non-differentially expressed genes within different TAD classes in the mutants. (A) Count and proportion of DEG and non-DEG in the three mutants (BEAF-32 knockdown, Cp190 Chro double knockdown and BEAF-32 Dref double knockdown) within TADs that: are fully conserved; lose one or both borders; have slightly shifted borders in the mutants. (B) We group different TAD classes in two subgroups (blue and yellow) and performed a permutation test to investigate if DEG overlap with any of the class more than expected by chance and plot the corresponding proportion of DEG over non-DEG for each group together with the corresponding -log_10_ p-values.

**Figure S7.**
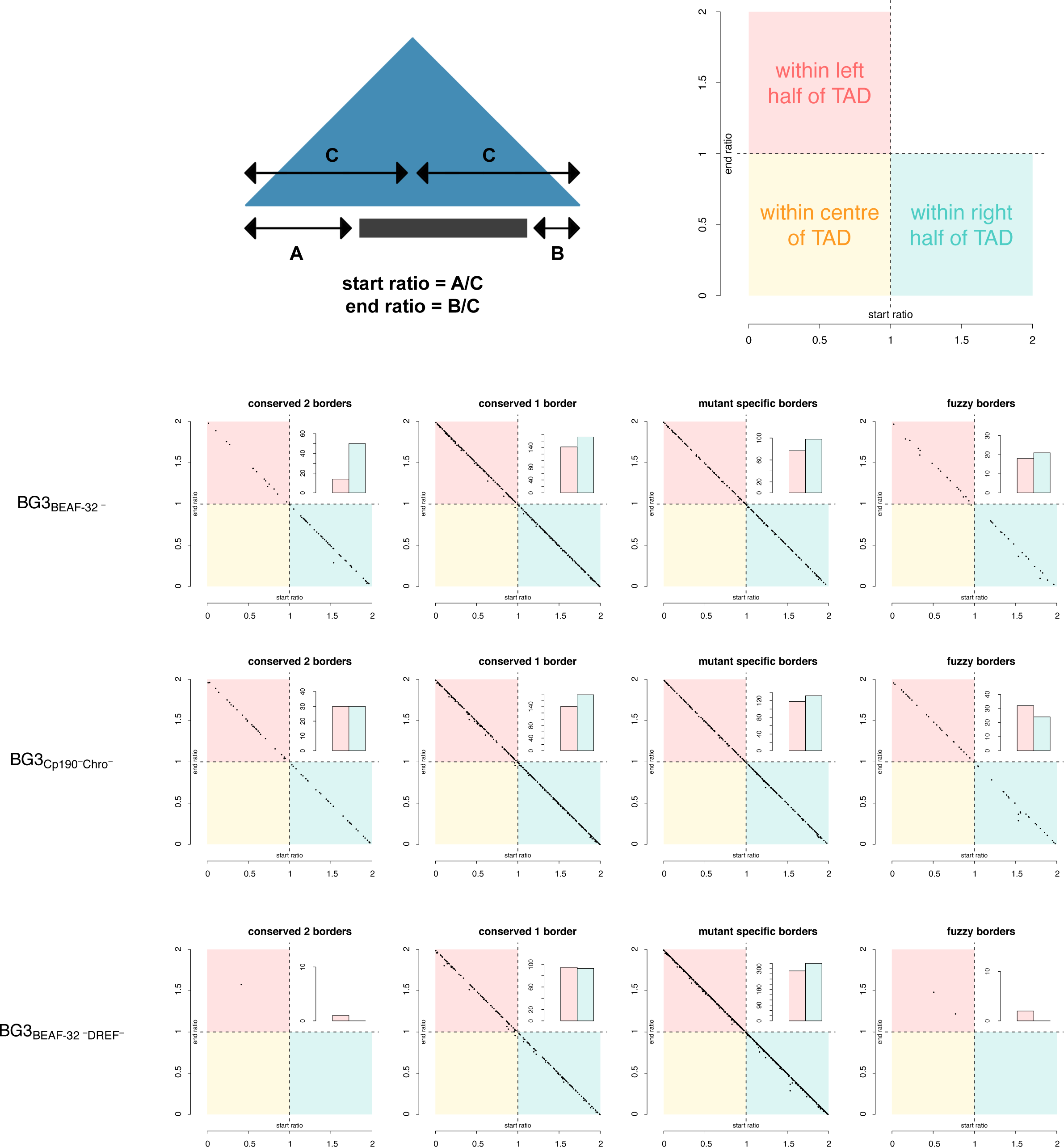
The allocation of differentially expressed genes within robust TADs. Allocation and count of differentially expressed genes within robust TADs in three mutants (BEAF-32 knockdown, Cp190 and Chro double knockdown, BEAF-32 Dref double knockdown) based on start and end ratios. The start ratio is defined as a distance from the left border of the TAD to the start position of the gene divided by the half of TAD size, where a start ratio bigger than 1 means that the gene is allocated on the right side of the TAD (blue square). The end ratio is defined as a distance from the right border of the TAD to the end position of the gene divided by the half of TAD size, where an end ratio bigger than 1 indicates the gene allocated on the left side of the TAD (red square). Genes having both start and end ratio less than 1 are allocated within TAD centre (yellow square). The majority of differentially expressed genes occupy less than half of the TAD. Only couple of genes are allocated within TAD centre – they are very close to point (1,1) indicating relatively short genes. Majority of genes are allocated either on the left or the right half of the TAD with no strong bias towards TAD borders.

**Figure S8.**
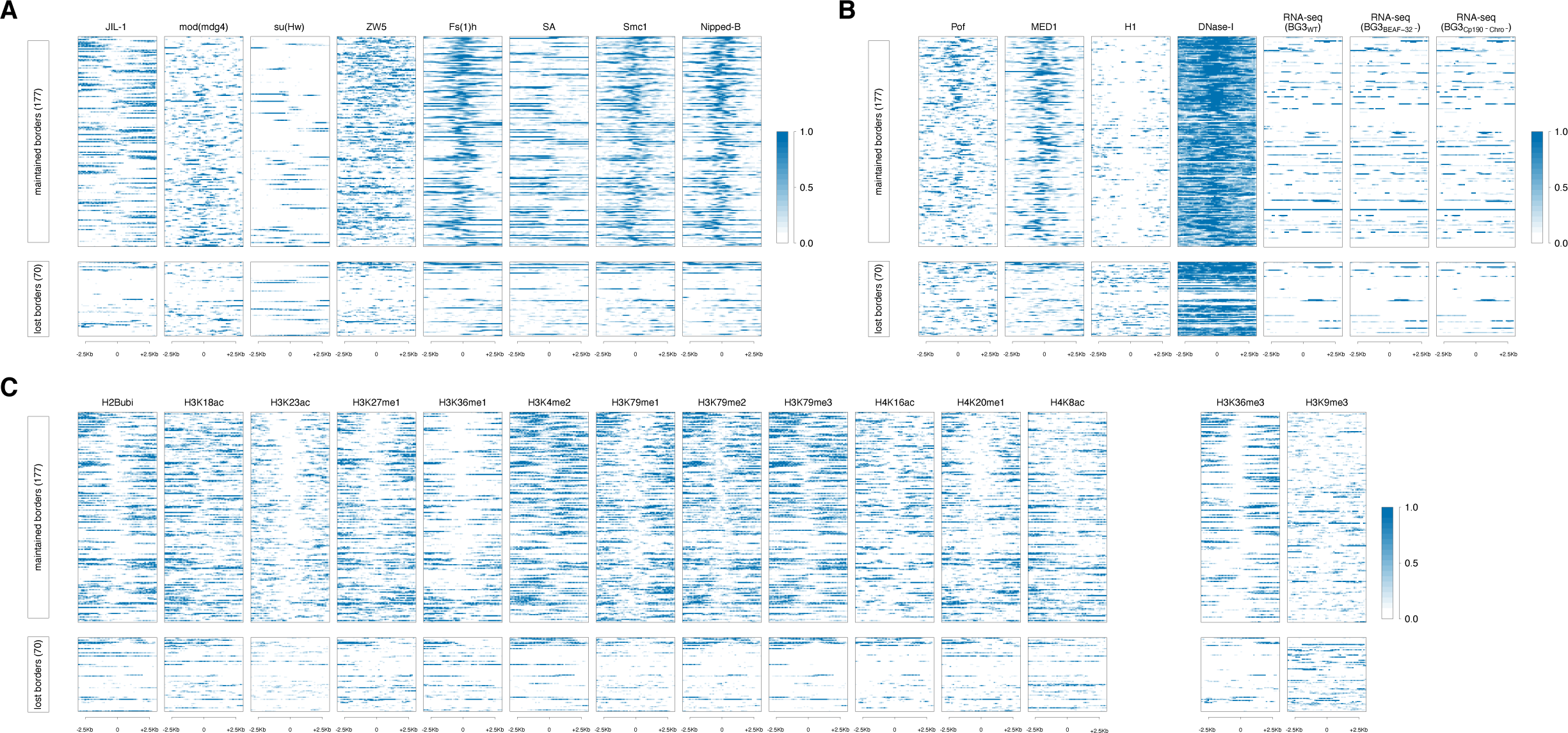
Additional chromatin features enrichment at TAD borders. (A) Profiles of architectural proteins (JIL-1, mod(mdg4), su(Hw), ZW5, Fs(1)h, SA, Smc1 and Nipped-B) around direct maintained and lost TAD borders. (B) Profiles of accessibility (H1 and DNase-I) and transcription (MED1 and Pof) at maintained and lost TAD borders. In addition, we plot RNA-seq signal in WT, BEAF-32 knockdown and Cp190 and Chro double knockdown. (C) Profiles of histone modifications (H2Bubi, H3K18ac, H3K23ac, H3K27me1, H3K36me1, H3K4me2, H3K79me1, H3K79me2, H3K79me3, H4K16ac, H4K20me1, H4K8ac, H3K36me3 and H3K9me3) at direct maintained and lost TAD borders.

**Figure S9.**
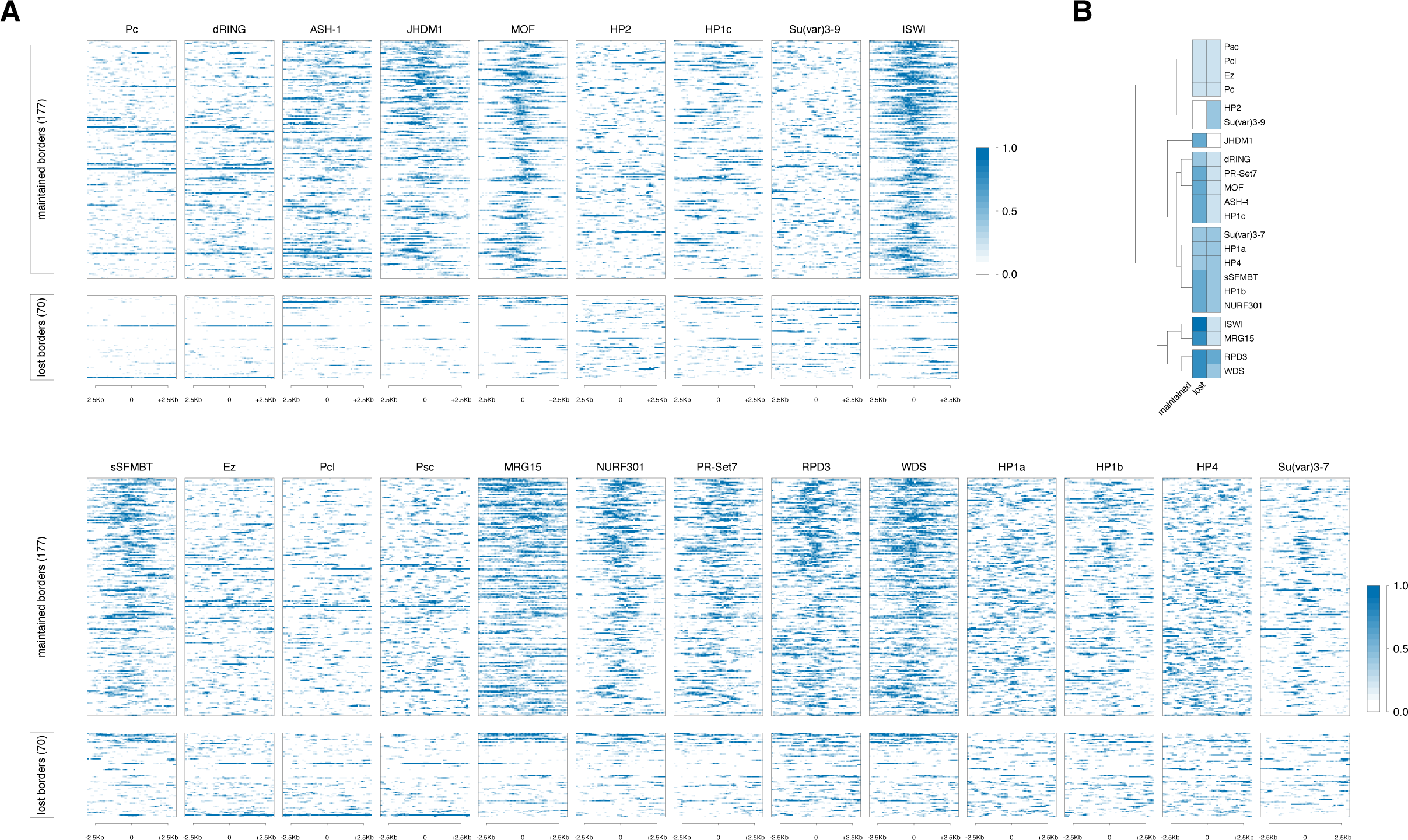
Enrichment of other proteins/complexes at TAD borders. (A) Profiles of Pc, dRING, ASH−1, JHDM1, MOF, HP2, HP1c, Su(var)3-9, ISWI, sSFMBT, Ez, Pcl, Psc, MRG15, NURF301, PR-Set7, RPD3, WDS, HP1a, HP1b, HP4 and Su(var)3-7 at direct maintained and lost TAD borders. (B) Clustering of the signal at direct maintained and lost TAD borders.

**Figure S10.**
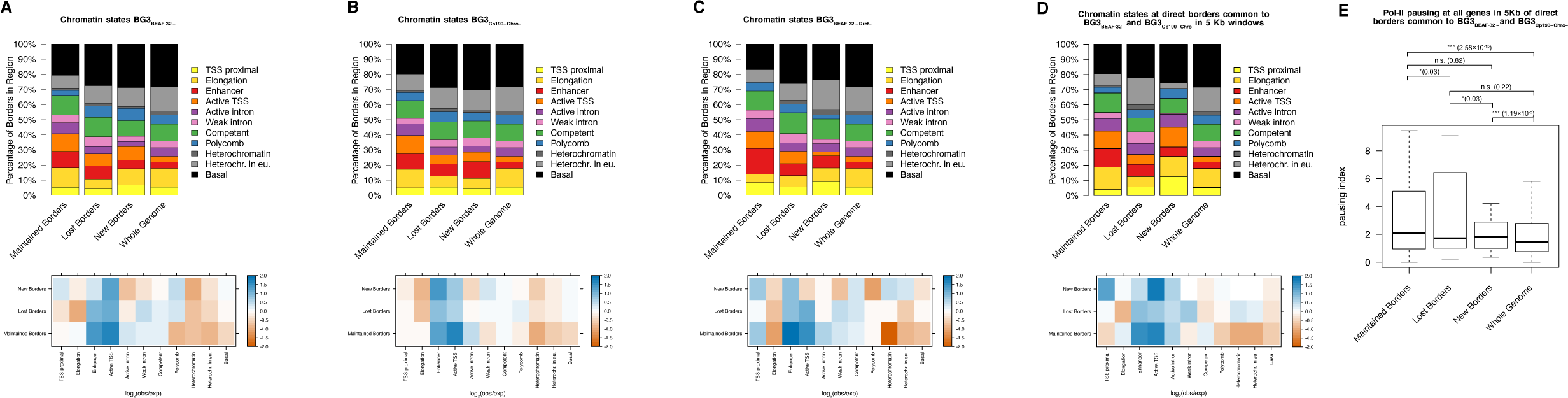
Chromatin state annotation at maintained, lost and new TAD borders. We computed the overlap between each maintained, lost and new robust TAD border (see Figures 1C and 2B) with chromatin states in the three mutants: (A) BEAF-32 single knockdown, (B) Cp190 and Chro double knockdown and (C) BEAF-32 and Dref double knockdown. Top panels: the percentage of the overlap between the different chromatin states and maintained, lost and new robust TAD borders and whole genome. Bottom panels: log_2_(observed/expected), where observed is the percentage in maintained, lost or new classes and expected is the whole genome distribution. (D) Same as in (A-C) for maintained, lost and new direct borders that were common between BEAF-32 single knockdown and Cp190 and Chro double knockdown mutants. (E) Pol-II pausing at maintained, lost and new direct borders that were common in BEAF-32 single knockdown and Cp190 and Chro double knockdown mutants. We performed a Mann-Whitney U test (p-value: n.s. ≥ 0.05, * p-value < 0.05, ** < 0.01 and *** < 0.001).

**Figure S11.**
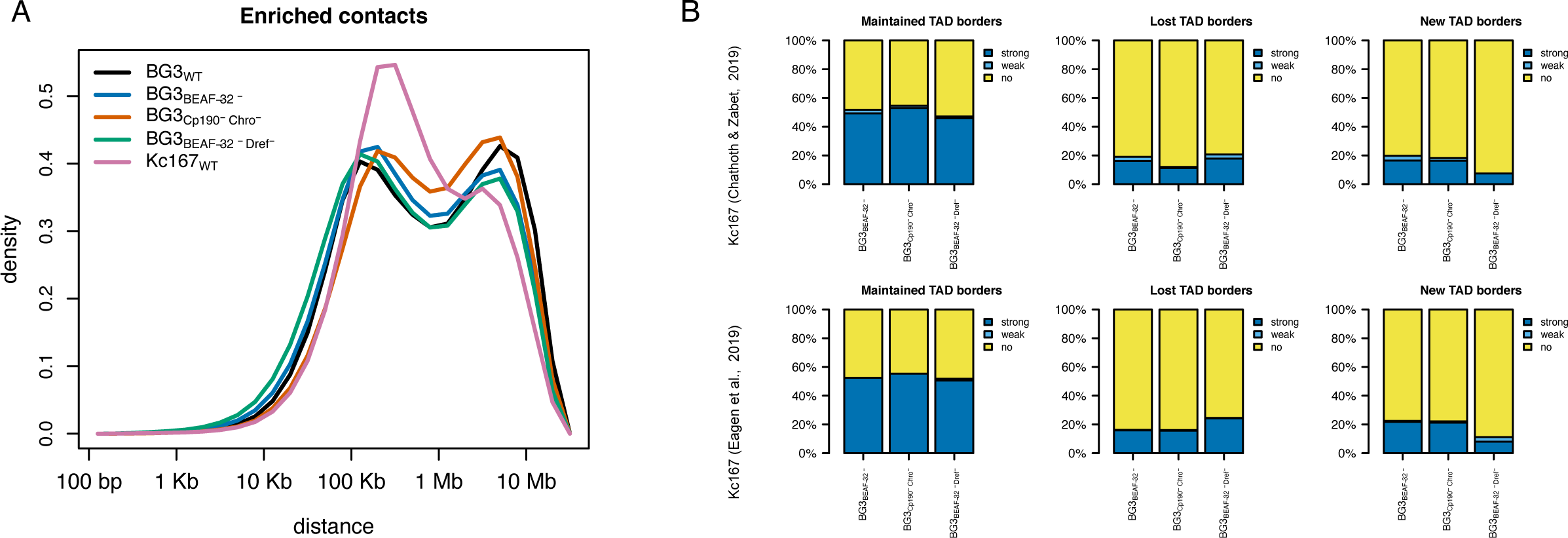
Comparison of the 3D chromatin organisation in the mutants with 3D chromatin organisation in Kc167 cells. (A) Density distribution of enriched number of contacts. We highlight the density plot of the enriched contacts instead of counts. This was necessary in order to take into account the differences generated by sequencing depth of the libraries for the different mutants. For Kc167 we used data from ^26^. (B) Proportion of robust TAD borders that are maintained in the three mutants and are also present in Kc167 cells as either strong or weak borders. Top panels: annotation of robust TAD borders in Kc167 using data from ^26^. Bottom panels: annotation of TAD borders in Kc167 using a very high sequencing depth library from ^48^.

**Figure S12.**
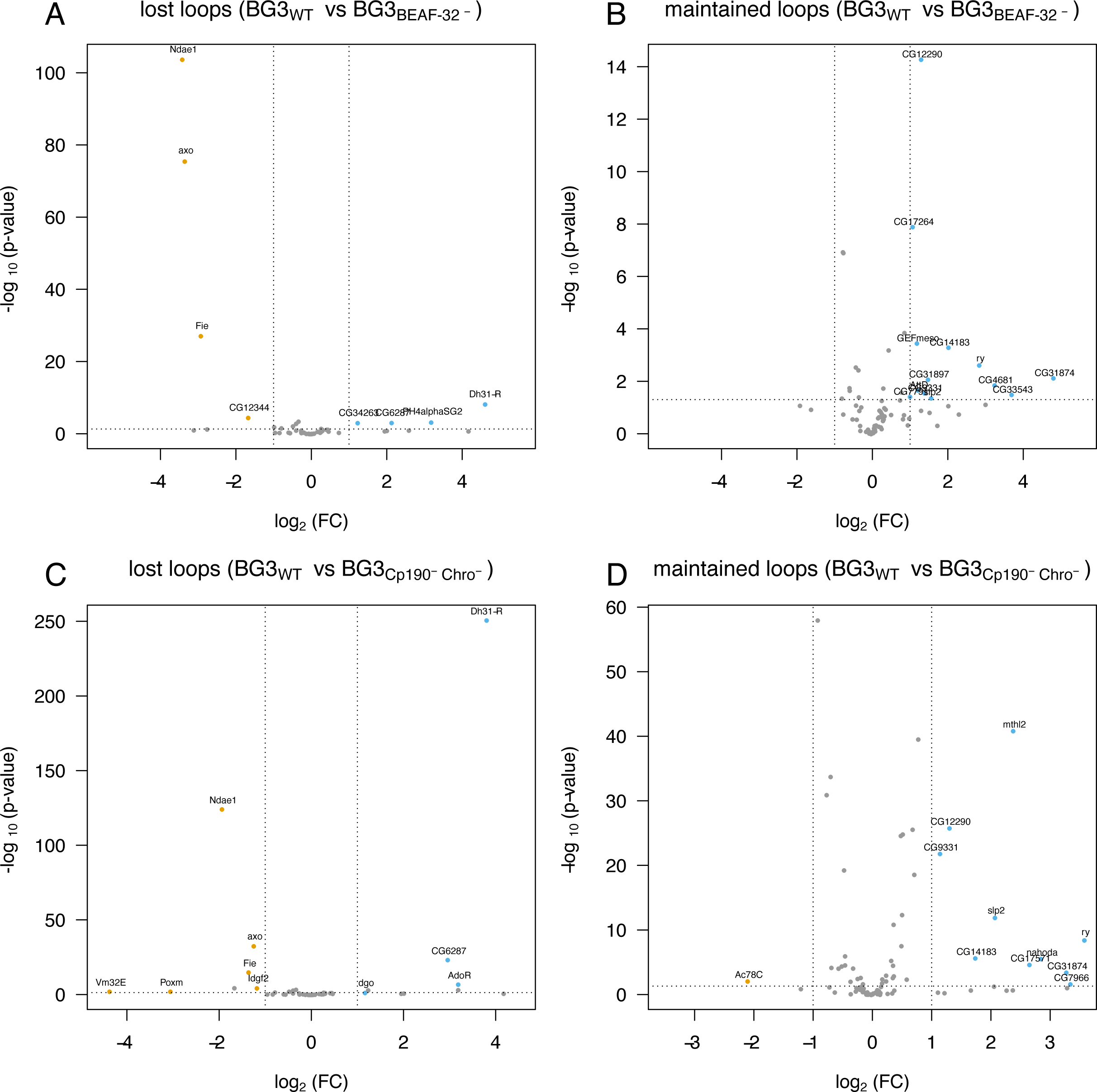
Differential expression of genes located at chromatin loops. Volcano plots of the -log_10_ of the adjusted p-value as a function of log_2_ fold change. We considered all genes located at lost chromatin loops (A and C) and maintained chromatin loops (B and D) in BEAF-32 single knockdown (A and B) and Cp190 and Chro double knockdown (C and D).

**Figure S13.**
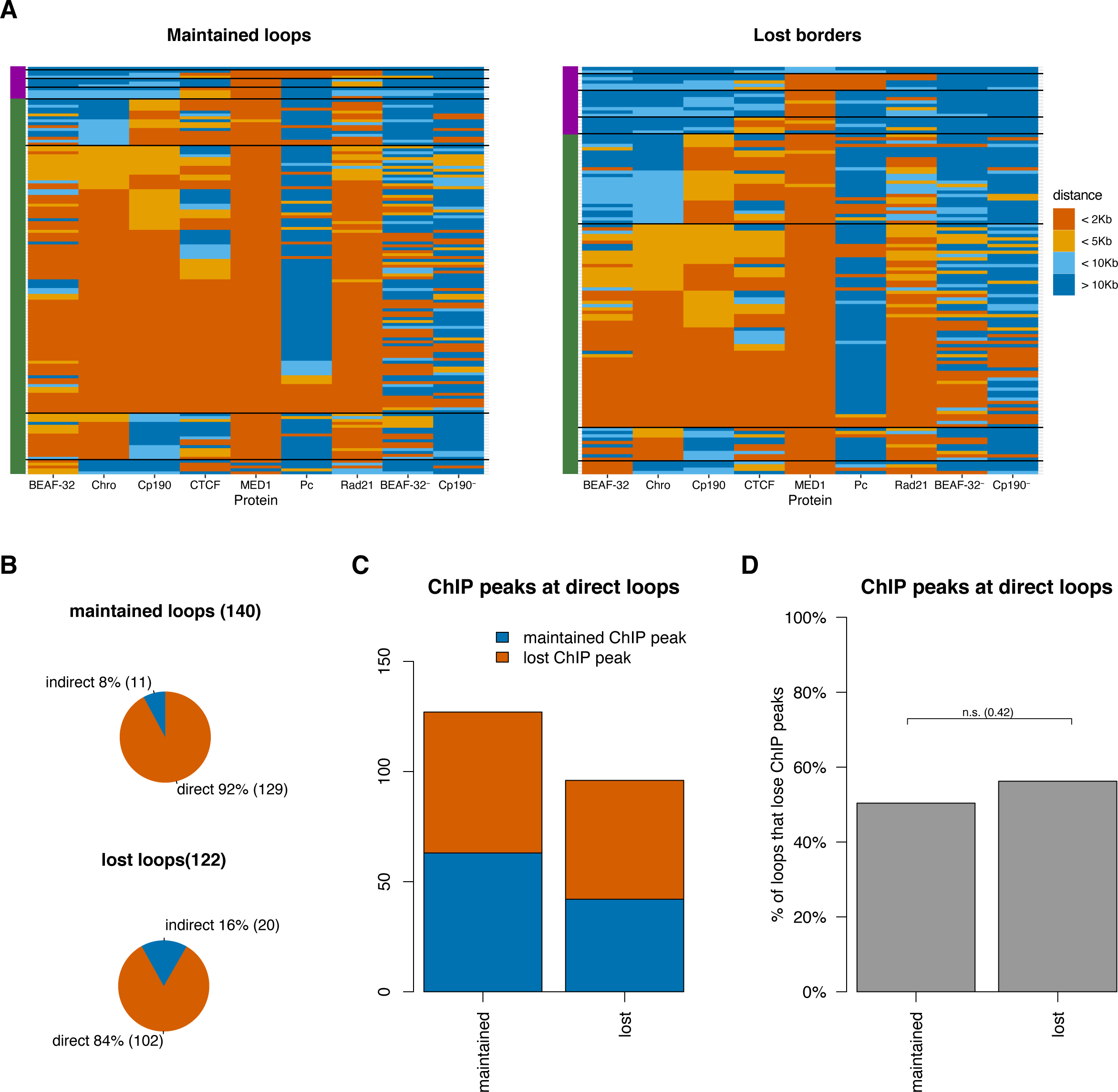
Direct and indirect chromatin loops. We considered separately the cases of maintained and lost chromatin loops. (A) Heatmaps plotting the distance of the closest ChIP peak from a maintained or lost loop for: BEAF-32 (WT and BEAF-32 knockdown), Chro (WT), Cp190 (WT and Cp190 knockdown), CTCF (WT), MED1 (WT), Pc (WT) and Rad21 (WT). Green bar on the side of each heatmap marks direct loops (loops that show binding of BEAF-32, Chro and/or Cp190), while purple indirect loops (all other borders). (B) Percentage of maintained and lost chromatin loops that have direct binding of BEAF-32, Cp190 and Chro. (C) number of chromatin loops that have BEAF-32 or Cp190 ChIP in WT cells and lose those peaks in BEAF-32 and Cp190 single knockdowns. (D) Percentage of chromatin loops that have BEAF-32 or Cp190 ChIP in WT and lose them in the in BEAF-32 and Cp190 single knockdowns. We performed a Fisher’s exact test and the corresponding p-value is displayed above the barplots.

**Figure S14.**
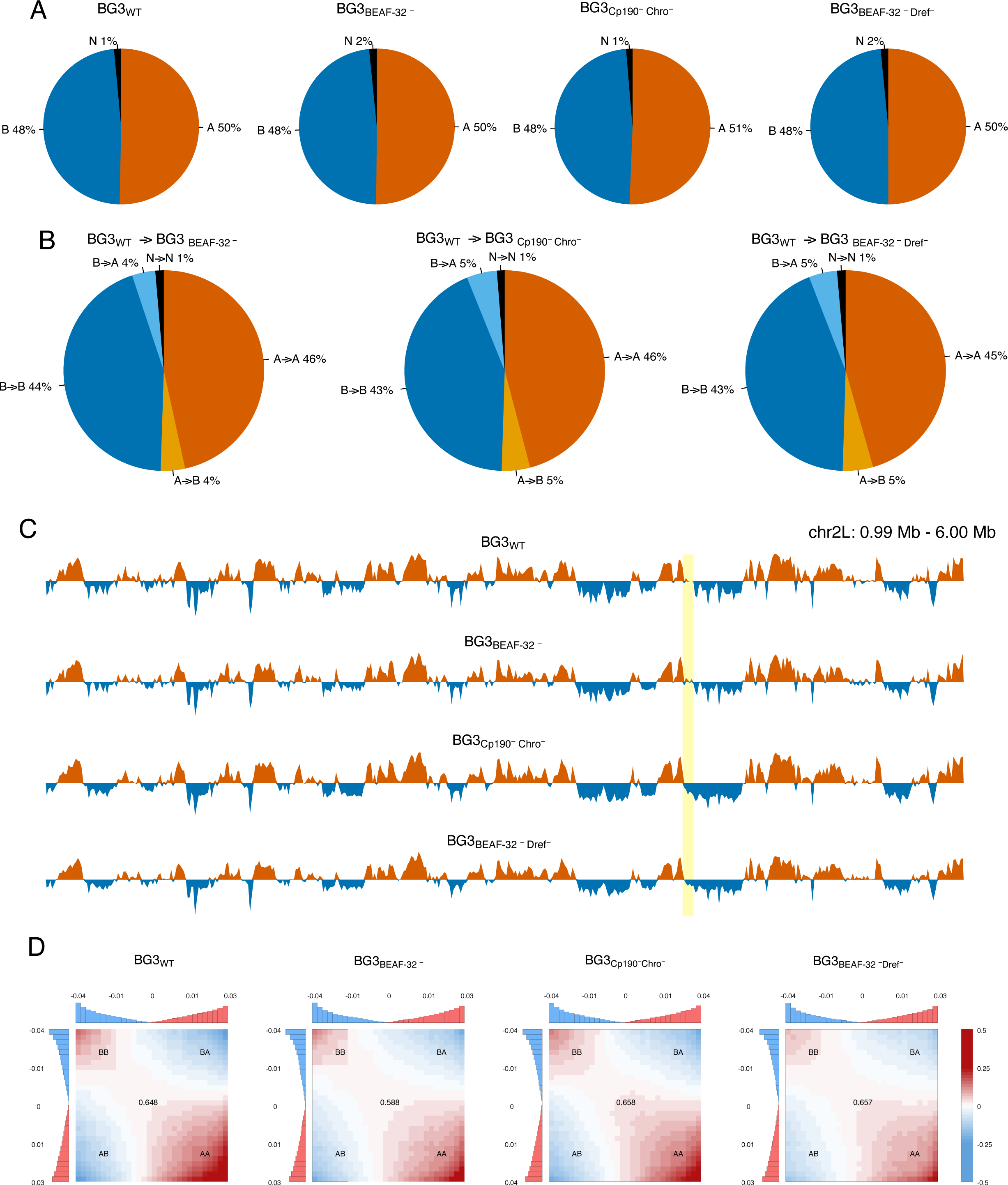
A/B compartments in BG3 WT and mutant cells. (A) Percentage of the genome that was called as A or a B compartment in WT cells, BEAF-32 knockdown cells, Cp190 and Chro double knockdown cells and BEAF-32 and Dref double knockdown cells. The compartments were detected in 10 Kb bins. Regions that could not be classified as either an A or a B compartment were labelled as N. (B) Percentage of A/B compartments that was maintained or switched in the three mutants. (C) Profile of eigen vectors on chromosome 2L. Red values indicated positive values (A compartment) and blue values indicate negative values (B compartment). One example of compartment switching is highlighted. (D) Averaged Pearson correlation profiles stratified by eigenvector percentile calculated as described in Methods. The top and left panels show averages over 30 percentile bins in absolute values: in blue there are partitions with negative average eigenvector scores (regions belonging to B compartments), in red there are partitions with positive average eigenvector scores (regions belonging to A compartments). Numbers at the centre of the heat maps indicate compartment strength calculated as the ratio of (AA+BB)/(AB+BA) using the mean values from corner sub-matrices of 10×10 bins.

**Figure S15.**
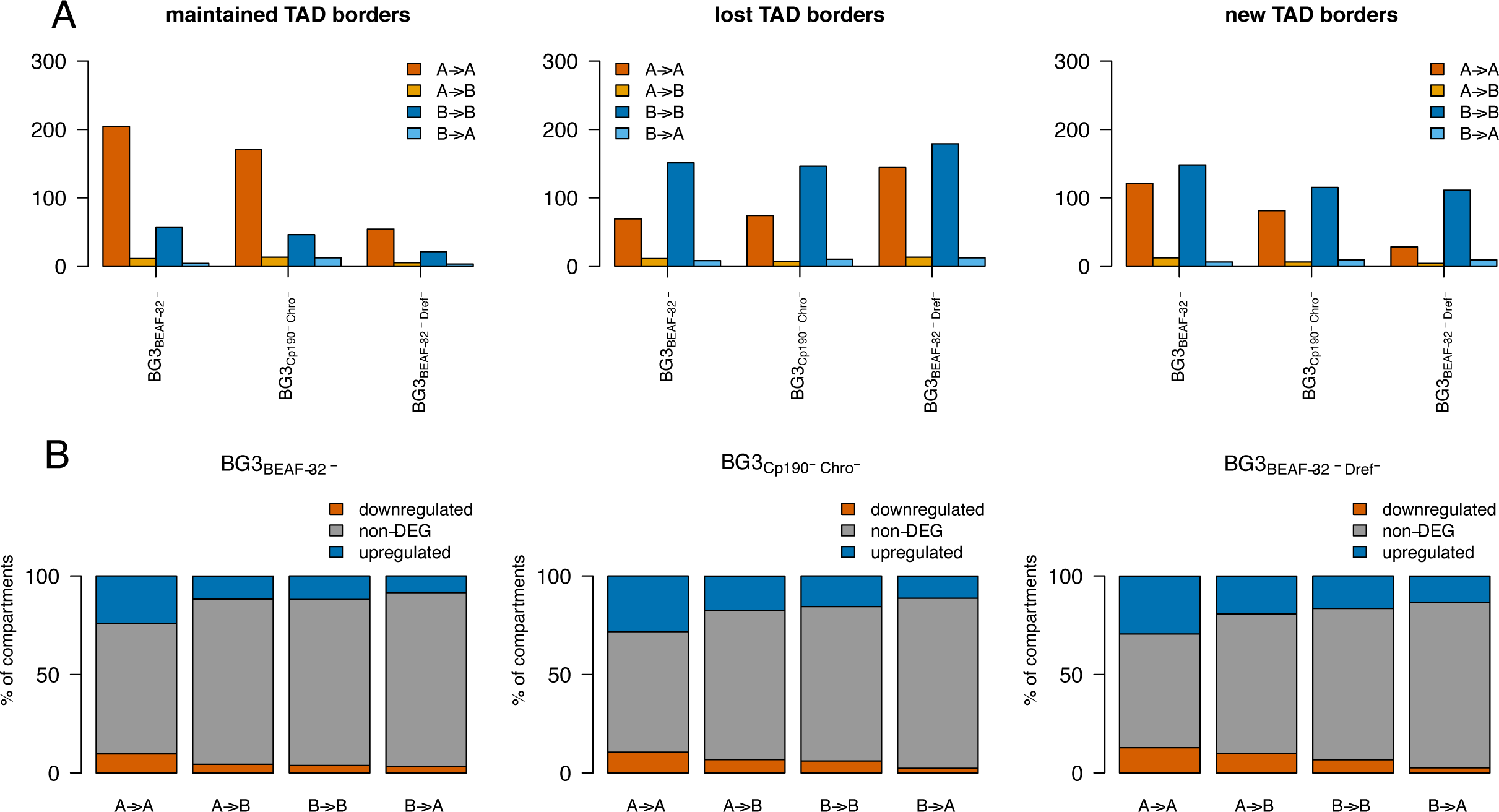
Characterisation of compartments based on TADs and differential gene expression. (A) Distribution of maintained, lost and new TAD borders in different classes of compartments switching for the three mutants. (B) Percentage of compartments switching that overlap with DEG and non-DEGs. We used a log_2_ fold change of 1.0 when computing DEGs. Majority of compartments switch do not have any DEG.

**Figure S16.**
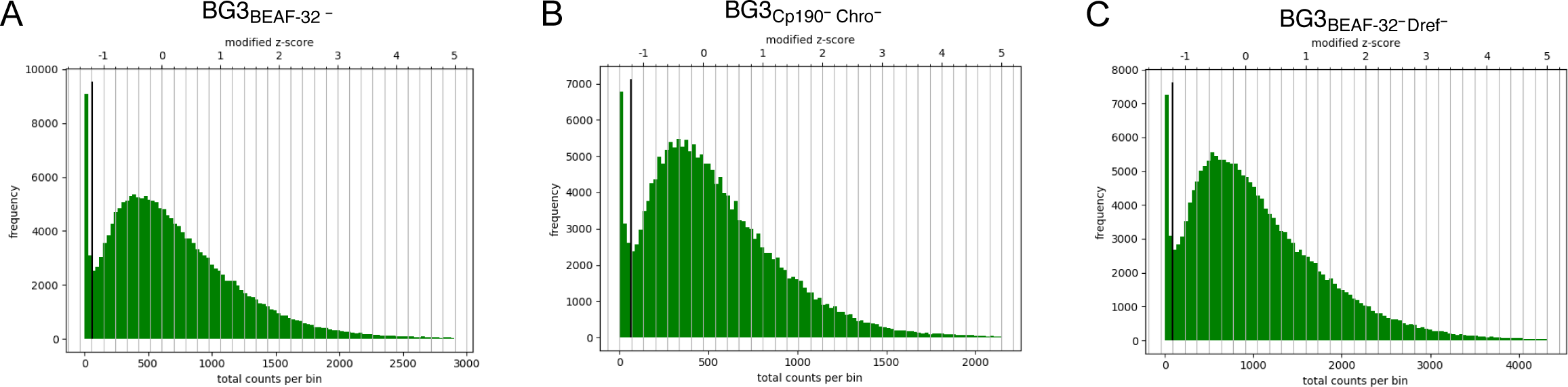
Diagnostic plots for correction of Hi-C plots from HiCExplorer. Histograms of the sum of contact per bin in (A) BEAF-32 single knockdown, (B) Cp190 Chro double knockdown and (C) BEAF-32 Dref double knockdown. The vertical black line represents the lower threshold for removing bins with lower number of reads.

## References

1. Dixon, J. R. et al. Topological domains in mammalian genomes identified by analysis of chromatin interactions. Nature 485, 376–380 (2012).

2. Sexton, T. et al. Three-dimensional folding and functional organization principles of the Drosophila genome. Cell 148, 458–72 (2012).

3. Vietri Rudan, M., et al. Comparative Hi-C Reveals that CTCF Underlies Evolution of Chromosomal Domain Architecture. Cell Rep. 10, 1297–1309 (2015).

4. Ghavi-Helm, Y. et al. Enhancer loops appear stable during development and are associated with paused polymerase. Nature 512, 96–100 (2014).

5. Dixon, J. R. et al. Chromatin architecture reorganization during stem cell differentiation. Nature 518, 331–336 (2015).

6. Li, L. et al. Widespread Rearrangement of 3D Chromatin Organization Underlies Polycomb-Mediated Stress-Induced Silencing. Mol. Cell 58, 216–231 (2015).

7. Lupianez, D. G. et al. Disruptions of Topological Chromatin Domains Cause Pathogenic Rewiring of Gene-Enhancer Interactions. Cell 161, 1012–1025 (2015).

8. Kraft, K. et al. Serial genomic inversions induce tissue-specific architectural stripes, gene misexpression and congenital malformations. Nat. Cell Biol. 21, 305–310 (2019).

9. Flavahan, W. A. et al. Insulator dysfunction and oncogene activation in IDH mutant gliomas. Nature 529, 110–114 (2016).

10. Hnisz, D. et al. Activation of proto-oncogenes by disruption of chromosome neighborhoods. Science (80-.). 351, 1454–1458 (2016).

11. Chandra, T. et al. Global Reorganization of the Nuclear Landscape in Senescent Cells. Cell Rep. 10, 471–483 (2015).

12. Sun, J. H. et al. Disease-Associated Short Tandem Repeats Co-localize with Chromatin Domain Boundaries. Cell 175, 224–238.e15 (2018).

13. Akdemir, K. C. et al. Disruption of chromatin folding domains by somatic genomic rearrangements in human cancer. Nat. Genet. 52, 294–305 (2020).

14. Cubenãs-Potts, C. et al. Different enhancer classes in Drosophilabind distinct architectural proteins and mediate unique chromatin interactions and 3D architecture. Nucleic Acids Res. 45, 1714–1730 (2017).

15. Harmston, N. et al. Topologically associated domains are ancient features that coincide with Metazoan clusters of extreme noncoding conservation. Nat. Commun. 8, 441 (2017).

16. Van Bortle, K. et al. Insulator function and topological domain border strength scale with architectural protein occupancy. Genome Biol. 15, R82 (2014).

17. Stadler, M. R., Haines, J. E. & Eisen, M. Convergence of topological domain boundaries, insulators, and polytene interbands revealed by high-resolution mapping of chromatin contacts in the early Drosophila melanogaster embryo. Elife 6, e29550 (2017).

18. Rao, S. S. P. et al. A 3D Map of the Human Genome at Kilobase Resolution Reveals Principles of Chromatin Looping. Cell 159, 1665–1680 (2014).

19. de Wit, E. TADs as the Caller Calls Them. Journal of Molecular Biology vol. 432 638–642 (2020).

20. Mirny, L. A., Imakaev, M. & Abdennur, N. Two major mechanisms of chromosome organization. Current Opinion in Cell Biology vol. 58 142–152 (2019).

21. Zuin, J. et al. Cohesin and CTCF differentially affect chromatin architecture and gene expression in human cells. Proc. Natl. Acad. Sci. U. S. A. 111, 996–1001 (2014).

22. Nora, E. P. et al. Targeted Degradation of CTCF Decouples Local Insulation of Chromosome Domains from Genomic Compartmentalization. Cell 169, 930–944.e22 (2017).

23. Schwarzer, W. et al. Two independent modes of chromatin organization revealed by cohesin removal. Nature 551, 51–56 (2017).

24. El-Sharnouby, S. et al. Regions of very low H3K27me3 partition the Drosophila genome into topological domains. PLoS One 12, 1–23 (2017).

25. Ramirez, F. et al. High-resolution TADs reveal DNA sequences underlying genome organization in flies. Nat. Commun. 9, 189 (2018).

26. Chathoth, K. T. & Zabet, N. R. Chromatin architecture reorganisation during neuronal cell differentiation in Drosophila genome. Genome Res. 29, 613–625 (2019).

27. Matthews, N. E. & White, R. Chromatin Architecture in the Fly: Living without CTCF/Cohesin Loop Extrusion? BioEssays 41, 1900048 (2019).

28. Rowley, M. J. et al. Condensin II Counteracts Cohesin and RNA Polymerase II in the Establishment of 3D Chromatin Organization. Cell Rep. 26, 2890–2903.e3 (2019).

29. Wang, Q., Sun, Q., Czajkowsky, D. M. & Shao, Z. Sub-kb Hi-C in D. melanogaster reveals conserved characteristics of TADs between insect and mammalian cells. Nat. Commun. 9, 188 (2018).

30. Martin, P. C. N. & Zabet, N. R. Dissecting the binding mechanisms of transcription factors to DNA using a statistical thermodynamics framework. bioRxiv (2019) doi:10.1101/666446.

31. Mathelier, A. et al. {JASPAR 2014}: an extensively expanded and updated open-access database of transcription factor binding profiles. Nucleic Acids Res. 42, D142–D147 (2014).

32. Hirose, F. et al. Isolation and characterization of cDNA for DREF, a promoter-activating factor for Drosophila DNA replication-related genes. J. Biol. Chem. 271, 3930–3937 (1996).

33. Vogelmann, J. et al. Chromatin Insulator Factors Involved in Long-Range {DNA} Interactions and Their Role in the Folding of the \emph{Drosophila} Genome. PLoS Genet 10, e1004544 (2014).

34. Ghavi-Helm, Y. et al. Highly rearranged chromosomes reveal uncoupling between genome topology and gene expression. Nat. Genet. 51, 1272–1282 (2019).

35. Williamson, I. et al. Developmentally regulated Shh expression is robust to TAD perturbations. Dev. 146, (2019).

36. Schwartz, Y. B. et al. Nature and function of insulator protein binding sites in the Drosophila genome. Genome Res. 22, 2188–2198 (2012).

37. Kharchenko, P. V et al. Comprehensive analysis of the chromatin landscape in Drosophila melanogaster. Nature (2010) doi:10.1038/nature09725.

38. Ulianov, S. V et al. Active chromatin and transcription play a key role in chromosome partitioning into topologically associating domains. Genome Resarch 26, 70–84 (2016).

39. Rowley, M. J. et al. Evolutionarily Conserved Principles Predict 3D Chromatin Organization. Mol. Cell 67, 837–852.e7 (2017).

40. Ogiyama, Y., Schuettengruber, B., Papadopoulos, G. L., Chang, J.-M. & Cavalli, G. Polycomb-Dependent Chromatin Looping Contributes to Gene Silencing during Drosophila Development. Mol. Cell 71, 73–88.e5 (2018).

41. Szabo, Q. et al. TADs are 3D structural units of higher-order chromosome organization in Drosophila. Sci. Adv. 4, (2018).

42. Kwon, S. Y., Grisan, V., Jang, B., Herbert, J. & Badenhorst, P. Genome-Wide Mapping Targets of the Metazoan Chromatin Remodeling Factor NURF Reveals Nucleosome Remodeling at Enhancers, Core Promoters and Gene Insulators. PLOS Genet. 12, e1005969 (2016).

43. Saha, P., Sowpati, D. T., Soujanya, M., Srivastava, I. & Mishra, R. K. Interplay of pericentromeric genome organization and chromatin landscape regulates the expression of Drosophila melanogaster heterochromatic genes. Epigenetics and Chromatin 13, 41 (2020).

44. Sexton, T. et al. Three-Dimensional Folding and Functional Organization Principles of the \emph{Drosophila}phila Genome. Cell 148, 458–472 (2012).

45. Hug, C. B., Grimaldi, A. G., Kruse, K. & Vaquerizas, J. M. Chromatin Architecture Emerges during Zygotic Genome Activation Independent of Transcription. Cell 169, 216–228.e19 (2017).

46. Ramírez, F. et al. High-resolution TADs reveal DNA sequences underlying genome organization in flies. Nat. Commun. 9, 189 (2018).

47. Skalska, L. et al. Chromatin signatures at Notch-regulated enhancers reveal large-scale changes in H3K56ac upon activation. EMBO J. 34, 1889–1904 (2015).

48. Eagen, K. P., Lieberman Aiden, E. & Kornberg, R. D. Polycomb-mediated chromatin loops revealed by a subkilobase-resolution chromatin interaction map. Proc. Natl. Acad. Sci. (2017) doi:10.1073/pnas.1701291114.

49. Noordermeer, D. et al. Temporal dynamics and developmental memory of 3D chromatin architecture at Hox gene loci. Elife 3, e02557 (2014).

50. Lieberman-Aiden, E. et al. Comprehensive Mapping of Long-Range Interactions Reveals Folding Principles of the Human Genome. Science (80-.). 326, 289–293 (2009).

51. Gibcus, J. H. et al. A pathway for mitotic chromosome formation. Science (80-.). 359, (2018).

52. Björkegren, C., Baranello, L., Björkegren, C. & Baranello, L. DNA Supercoiling, Topoisomerases, and Cohesin: Partners in Regulating Chromatin Architecture? Int. J. Mol. Sci. 19, 884 (2018).

53. Benedetti, F., Racko, D., Dorier, J., Burnier, Y. & Stasiak, A. Transcription-induced supercoiling explains formation of self-interacting chromatin domains in S. pombe. Nucleic Acids Res. 45, 9850–9859 (2017).

54. Mateo, L. J. et al. Visualizing DNA folding and RNA in embryos at single-cell resolution. Nature 568, 49–54 (2019).

55. Arzate-Mejía, R. G., Josué Cerecedo-Castillo, A., Guerrero, G., Furlan-Magaril, M. & Recillas-Targa, F. In situ dissection of domain boundaries affect genome topology and gene transcription in Drosophila. Nat. Commun. 11, 1–16 (2020).

56. Schwartz, Y. B. et al. Nature and function of insulator protein binding sites in the Drosophila genome. Genome Resarch 22, 2188–2198 (2012).

57. Bonev, B. et al. Multiscale 3D Genome Rewiring during Mouse Neural Development. Cell 171, 557–572.e24 (2017).

58. Rhodes, J. D. P. et al. Cohesin Disrupts Polycomb-Dependent Chromosome Interactions in Embryonic Stem Cells. Cell Rep. 30, 820–835.e10 (2020).

59. Muerdter, F. et al. Resolving systematic errors in widely used enhancer activity assays in human cells. Nat. Methods 15, 141–149 (2017).

60. Sanyal, A., Lajoie, B. R., Jain, G. & Dekker, J. The long-range interaction landscape of gene promoters. Nature 489, 109–113 (2012).

61. Javierre, B. M. et al. Lineage-Specific Genome Architecture Links Enhancers and Non-coding Disease Variants to Target Gene Promoters. Cell 167, 1369–1384.e19 (2016).

62. Zheng, M. et al. Multiplex chromatin interactions with single-molecule precision. Nature 566, 558–562 (2019).

63. Atlasi, Y. et al. Epigenetic modulation of a hardwired 3D chromatin landscape in two naive states of pluripotency. Nat. Cell Biol. 21, 568–578 (2019).

64. Espinola, S. M. et al. Cis -regulatory chromatin loops arise before TADs and gene activation, and are independent of cell fate during development. bioRxiv 2020.07.07.191015 (2020) doi:10.1101/2020.07.07.191015.

65. Ing-Simmons, E., Vaid, R., Mannervik, M. & Vaquerizas, J. Independence of 3D chromatin conformation and gene regulation during Drosophila dorsoventral patterning. bioRxiv 2020.07.07.186791 (2020) doi:10.1101/2020.07.07.186791.

66. Blanton, J., Gaszner, M. & Schedl, P. Protein:protein interactions and the pairing of boundary elements in vivo. Genes Dev. 17, 664–675 (2003).

67. Rath, U. et al. Chromator, a novel and essential chromodomain protein interacts directly with the putative spindle matrix protein skeletor. J. Cell. Biochem. 93, 1033–1047 (2004).

68. Whitfield, W. G., Millar, S. E., Saumweber, H., Frasch, M. & Glover, D. M. Cloning of a gene encoding an antigen associated with the centrosome in Drosophila. J. Cell Sci. 89, (1988).

69. Adams, M. D. et al. The Genome Sequence of Drosophila melanogaster. Science (80-). 287, 2185–2195 (2000).

70. dos Santos, G. et al. FlyBase: introduction of the Drosophila melanogaster Release 6 reference genome assembly and large-scale migration of genome annotations. Nucleic Acids Res. 43, D690–D697 (2015).

71. Li, H. & Durbin, R. Fast and accurate long-read alignment with Burrows-Wheeler transform. Bioinformatics 26, 589–595 (2010).

72. Durand, N. C. et al. Juicer Provides a One-Click System for Analyzing Loop-Resolution Hi-C Experiments. Cell Syst. 3, 95–98 (2017).

73. Imakaev, M. et al. Iterative correction of {HI-C} data reveals hallmarks of chromosome organization. Nat. Methods 9, 999–1003 (2012).

74. Naumova, N. et al. Organization of the mitotic chromosome. Science (80-.). 342, 948–953 (2013).

75. Abramo, K. et al. A chromosome folding intermediate at the condensin-to-cohesin transition during telophase. Nat. Cell Biol. 21, 1393–1402 (2019).

76. Brown, J. B. et al. Diversity and dynamics of the \emph{Drosophila} transcriptome. Nature (2014) doi:10.1038/nature12962.

77. Corrales, M. et al. Clustering of Drosophila housekeeping promoters facilitates their expression. Genome Res. 27, 1153–1161 (2017).

78. Bolger, A. M., Lohse, M. & Usadel, B. Trimmomatic: a flexible trimmer for Illumina sequence data. Bioinformatics 30, 2114–2120 (2014).

79. Kim, D. et al. TopHat2: accurate alignment of transcriptomes in the presence of insertions, deletions and gene fusions. Genome Biol. 14, R36 (2013).

80. Langmead, B. & Salzberg, S. L. Fast gapped-read alignment with {Bowtie} 2. Nat. Methods 9, 357–359 (2012).

81. Anders, S., Pyl, P. T. & Huber, W. HTSeq-A Python framework to work with high-throughput sequencing data. Bioinformatics 31, 166–169 (2015).

82. Love, M. I., Huber, W. & Anders, S. Moderated estimation of fold change and dispersion for RNA-seq data with DESeq2. Genome Biol. 15, 550 (2014).

83. Pherson, M., Misulovin, Z., Gause, M. & Dorsett, D. Cohesin occupancy and composition at enhancers and promoters are linked to DNA replication origin proximity in Drosophila. Genome Res. 29, 602–612 (2019).

84. Zhang, Y. et al. Model-based Analysis of {ChIP-Seq (MACS)}. Genome Biol. 9, R137 (2008).

85. Pherson, M. et al. Polycomb repressive complex 1 modifies transcription of active genes. Sci. Adv. 3, (2017).

86. Yanez-Cuna, J. O. et al. Dissection of thousands of cell type-specific enhancers identifies dinucleotide repeat motifs as general enhancer features. Genome Res. (2014) doi:10.1101/gr.169243.113.

87. Lun, A. T. L. & Smyth, G. K. diffHic: A Bioconductor package to detect differential genomic interactions in Hi-C data. BMC Bioinformatics 16, 258 (2015).

