## Supplementary Material for "The role of insulators and transcription in 3D chromatin organisation of flies"

**Table S1:** *Metrics from analysis of Hi-C library sequencing*

| Sample | Library size | Source | Total reads | mappable, unique and high quality | Pairs used | Inter-chromosomal | short range (<20kb) | long range |
|---|---|---|---|---|---|---|---|---|
| $BG_{WT}$ replicate 1 | 100% | GSE122603 | 151,493,083 | 101,206,098 | 42,160,814 | 7,157,538 | 12,969,585 | 22,033,691 |
| $BG_{WT}$ replicate 2 | 100% | GSE122603 | 147,730,225 | 95,703,042 | 42,999,219 | 7,851,274 | 13,180,675 | 21,967,270 |
| $BG_{BEAF-32^-}$ replicate 1 | 100% | this study | 172,597,536 | 69,859,110 | 32,240,219 | 4,463,615 | 9,357,013 | 18,419,591 |
| $BG_{BEAF-32^-}$ replicate 2 | 100% | this study | 156,655,569 | 81,740,811 | 41,458,552 | 5,690,882 | 11,839,584 | 23,928,086 |
| $BG_{Cp190^- Chro^-}$ replicate 1 | 100% | this study | 214,736,070 | 146,124,131 | 65,278,295 | 8,846,046 | 18,151,528 | 38,280,721 |
| $BG_{Cp190^- Chro^-}$ replicate 2 | 100% | this study | 167,983,753 | 104,046,976 | 45,830,412 | 6,142,658 | 13,111,506 | 26,576,248 |
| $BG_{BEAF-32^- Dref^-}$ replicate 1 | 100% | this study | 197,221,569 | 107,472,262 | 37,738,037 | 5,085,390 | 9,902,103 | 22,750,544 |
| $BG_{BEAF-32^- Dref^-}$ replicate 2 | 100% | this study | 190,654,840 | 53,234,566 | 18,645,676 | 2,462,801 | 5,210,579 | 10,972,296 |
| $Kc167_{WT}$ replicate 1 | 100% | GSE122603 | 152,371,706 | 98,701,754 | 49,712,803 | 3,532,342 | 16,386,181 | 29,794,280 |
| $Kc167_{WT}$ replicate 2 | 100% | GSE122603 | 188,531,576 | 117,170,016 | 61,211,572 | 4,522,842 | 17,549,275 | 39,139,455 |
| $BG_{WT}$ replicate 1 - 80% | 80% | GSE122603 | 121,195,638 | 81,045,523 | 34,528,646 | 5,857,796 | 10,713,800 | 17,957,050 |
| $BG_{WT}$ replicate 2 - 80% | 80% | GSE122603 | 118,173,473 | 76,656,728 | 35,462,491 | 6,480,416 | 10,908,557 | 18,073,518 |
| $BG_{BEAF-32^-}$ replicate 1 - 80% | 80% | this study | 138,073,566 | 55,898,357 | 26,805,854 | 3,703,534 | 7,833,118 | 15,269,202 |
| $BG_{BEAF-32^-}$ replicate 2 - 80% | 80% | this study | 125,326,239 | 65,443,211 | 33,860,610 | 4,647,796 | 9,764,035 | 19,448,779 |
| $BG_{Cp190^- Chro^-}$ replicate 1 - 80% | 80% | this study | 171,799,624 | 117,064,857 | 53,376,413 | 7,234,167 | 14,861,363 | 31,280,883 |
| $BG_{Cp190^- Chro^-}$ replicate 2 - 80% | 80% | this study | 134,385,744 | 83,338,488 | 37,245,423 | 4,993,379 | 10,680,858 | 21,571,186 |
| $BG_{BEAF-32^- Dref^-}$ replicate 1 - 80% | 80% | this study | 157,771,608 | 86,029,067 | 31,775,241 | 4,278,328 | 8,338,702 | 19,158,211 |
| $BG_{BEAF-32^- Dref^-}$ replicate 2 - 80% | 80% | this study | 152,530,267 | 42,612,149 | 15,651,046 | 2,066,595 | 4,379,973 | 9,204,478 |
| $Kc167_{WT}$ replicate 1 - 80% | 80% | GSE122603 | 121,887,879 | 78,989,128 | 40,649,289 | 2,888,640 | 13,471,045 | 24,289,604 |
| $Kc167_{WT}$ replicate 2 - 80% | 80% | GSE122603 | 150,825,628 | 93,798,562 | 50,505,474 | 3,752,258 | 14,525,871 | 32,227,345 |

**Table S2:** *Parameters for the correction of Hi-C matrices and the number of TADs*

| Sample | Library Size | Threshold DpnII | Threshold 10 Kb | Threhsold 100 Kb | Number of TAD borders | Number of strong TAD border |
|---|---|---|---|---|---|---|
| $BG_{WT}$ replicate 1 | 100% | [-1.4, 5] | [-1.4, 5] | [-2.4, 5] | | |
| $BG_{WT}$ replicate 2 | 100% | [-1.4, 5] | [-1.4, 5] | [-2.4, 5] | | |
| $BG_{WT}$ merged | 100% | [-1.4, 5] | [-1.4, 5] | [-2.4, 5] | 2260 | 989 |
| $BG_{BEAF-32^-}$ replicate 1 | 100% | [-1.0, 5] | [-0.8, 5] | [-1.9, 5] | | |
| $BG_{BEAF-32^-}$ replicate 2 | 100% | [-1.2, 5] | [-1.4, 5] | [-2.2, 5] | | |
| $BG_{BEAF-32^-}$ merged | 100% | [-1.2, 5] | [-1.4, 5] | [-2.2, 5] | 2557 | 1136 |
| $BG_{Cp190^- Chro^-}$ replicate 1 | 100% | [-1.2, 5] | [-1.4, 5] | [-2.4, 5] | | |
| $BG_{Cp190^- Chro^-}$ replicate 2 | 100% | [-1.2, 5] | [-1.4, 5] | [-2.4, 5] | | |
| $BG_{Cp190^- Chro^-}$ merged | 100% | [-1.2, 5] | [-1.4, 5] | [-2.4, 5] | 2163 | 869 |

| | | | | | | |
|---|---|---|---|---|---|---|
| BG$_{BEAF\text{-}32^{-}\ Dref^{-}}$ replicate 1 | 100% | [-1.2, 5] | [-1.4, 5] | [-2.4, 5] | | |
| BG$_{BEAF\text{-}32^{-}\ Dref^{-}}$ replicate 2 | 100% | [-1.2, 5] | [-1.4, 5] | [-2.4, 5] | | |
| BG$_{BEAF\text{-}32^{-}\ Dref^{-}}$ merged | 100% | [-1.2, 5] | [-1.4, 5] | [-2.4, 5] | 1417 | 441 |
| Kc167$_{WT}$ replicate 1 | 100% | [-1.4, 5] | [-1.4, 5] | [-2.6, 5] | | |
| Kc167$_{WT}$ replicate 2 | 100% | [-1.4, 5] | [-2.0, 5] | [-3.0, 5] | | |
| Kc167$_{WT}$ merged | 100% | [-1.4, 5] | [-1.6, 5] | [-3.0, 5] | 2512 | 1306 |
| BG$_{WT}$ replicate 1 - 80% | 80 % | | | | | |
| BG$_{WT}$ replicate 2 - 80% | 80 % | | | | | |
| BG$_{WT}$ merged - 80% | 80 % | [-1.4, 5] | | | 2179 | 902 |
| BG$_{BEAF\text{-}32^{-}}$ replicate 1 - 80% | 80 % | | | | | |
| BG$_{BEAF\text{-}32^{-}}$ replicate 2 - 80% | 80 % | | | | | |
| BG$_{BEAF\text{-}32^{-}}$ merged - 80% | 80 % | [-1.2, 5] | | | 2493 | 1074 |
| BG$_{Cp190^{-}\ Chro^{-}}$ replicate 1 - 80% | 80 % | | | | | |
| BG$_{Cp190^{-}\ Chro^{-}}$ replicate 2 - 80% | 80 % | | | | | |
| BG$_{Cp190^{-}\ Chro^{-}}$ merged - 80% | 80 % | [-1.2, 5] | | | 2051 | 776 |
| BG$_{BEAF\text{-}32^{-}\ Dref^{-}}$ replicate 1 - 80% | 80 % | | | | | |
| BG$_{BEAF\text{-}32^{-}\ Dref^{-}}$ replicate 2 - 80% | 80 % | | | | | |
| BG$_{BEAF\text{-}32^{-}\ Dref^{-}}$ merged - 80% | 80 % | [-1.2, 5] | | | 1346 | 416 |
| Kc167$_{WT}$ replicate 1 - 80% | 80 % | | | | | |
| Kc167$_{WT}$ replicate 2 - 80% | 80 % | | | | | |
| Kc167$_{WT}$ merged - 80% | 80 % | [-1.4, 5] | | | 2479 | 1229 |

**Table S3:** *Metrics from analysis of RNA-seq library sequencing*

| Sample | Total reads | After trimming | Aligned pairs | Concordant alignment rate | Multiple alignments | DEG$_{(log_2FC=2.0)}$ |
|---|---|---|---|---|---|---|
| BG$_{WT}$ replicate 1 | 24,486,765 | 23,334,081 | 19,022,098 | 80.2% | 15.9% | |
| BG$_{WT}$ replicate 2 | 24,953,334 | 23,902,047 | 19,891,746 | 82.0% | 14.9% | |
| BG$_{WT}$ replicate 3 | 25,765,693 | 24,761,665 | 20,804,226 | 82.4% | 15.3% | |
| BG$_{BEAF\text{-}32^{-}}$ replicate 1 | 22,783,325 | 21,691,464 | 17,512,866 | 78.8% | 20.5% | |
| BG$_{BEAF\text{-}32^{-}}$ replicate 2 | 26,110,498 | 25,485,051 | 21,761,711 | 83.6% | 20.1% | 596 |
| BG$_{BEAF\text{-}32^{-}}$ replicate 3 | 22,390,841 | 21,402,262 | 17,408,320 | 78.5% | 29.1% | |
| BG$_{Cp190^{-}\ Chro^{-}}$ replicate 1 | 29,595,777 | 27,953,418 | 22,271,222 | 77.6% | 22.5% | |
| BG$_{Cp190^{-}\ Chro^{-}}$ replicate 2 | 25,622,361 | 24,395,295 | 19,674,867 | 78.6% | 21.8% | 687 |
| BG$_{Cp190^{-}\ Chro^{-}}$ replicate 3 | 33,314,606 | 32,443,121 | 27,496,068 | 82.9% | 21.6% | |
| BG$_{BEAF\text{-}32^{-}\ Dref^{-}}$ replicate 1 | 30,470,304 | 29,551,901 | 25,119,383 | 83.1% | 23.8% | |

| | | | | | | |
|---|---|---|---|---|---|---|
| $0BG_{BEAF\text{-}32^{-}\ Dref^{-}}$ replicate 2 | 21,519,093 | 20,564,923 | 17,119,372 | 81.2% | 24.2% | 810 |
| $BG_{BEAF\text{-}32^{-}\ Dref^{-}}$ replicate 3 | 25,156,762 | 23,927,406 | 19,266,708 | 77.9% | 25.0% | |

**Table S4:** *Classification of genes located at maintained TAD borders*. We considered all maintained TAD borders that are common between BEAF-32 single mutant and Cp190 and Chro double mutant and that are also present in Kc167 cells (159 borders). We selected all genes within 5 Kb (257) and then identified their expression levels in 85 tissues/developmental times or cell lines (252 genes). Genes that were expressed in 40th percentile of expression in all 85 samples were classified as house keeping genes.

**Table S5:** *Primer sequences used for RNAi and RT-qPCR.*

| | |
|---|---|
| **RNAi primer sequences** | |
| BEAF-32_F(RNAi) | TAATACGACTCACTATAGGGGGAGGAGTACGAGCAGAACG |
| BEAF-32_R(RNAi) | TAATACGACTCACTATAGGGACGCTGATTTGCCCATTTAC |
| Chro_F(RNAi) | TAATACGACTCACTATAGGGCTTTGTTATTCGCACAGGCA |
| Chro_R(RNAi) | TAATACGACTCACTATAGGGCAGGAGGAATTGGCAAACAT |
| Cp190_F(RNAi) | TAATACGACTCACTATAGGGCGGCATGGACATCATCATAA |
| Cp190_R(RNAi) | TAATACGACTCACTATAGGGACAGTTGGACAGCCCAGTTC |
| Dref_F(RNAi) | TAATACGACTCACTATAGGGCGAGATACCAAATCCTCCGA |
| Dref_R(RNAi) | TAATACGACTCACTATAGGGTCGCCAGTGCAGACTAATTG |
| **RT-qPCR primer sequences** | |
| BEAF-32_F | AGGATCCACTGTGCTATAGTCC |
| BEAF-32_R | GCTGGTGAAGTCGAATGGGT |
| Chro_F | AGTTTAAAGCTATCGACAGG |
| Chro_R | CAGAGATGATTTGGTTCCG |
| Cp190_F | CACCGACTACTTCAATGTAC |
| Cp190_R | TTTTAAGCTCAAACTCCAGG |
| Dref_F | CCCAAGATGAAAAGCGTATA |
| Dref_R | CCTTGTGACACTTAATGCAGA |
| RpL32_F | AAGCGGCGACGCACTCTGTT |
| RpL32_R | GCCCAGCATACAGGCCCAAG |

**Table S6:** *Datasets for architectural proteins used in this work*

| Architectural proteins | | | dm3 or dm6 | LiftOver to dm6 |
|---|---|---|---|---|
| BEAF-32 | 921 | GSE20811 | dm3 | yes |
| CTCF | 3673 | GSE32783 | dm3 | yes |
| CP190 | 924 | GSE20814 | dm3 | yes |
| Criz/Chrom | 275 | GSE20761 | dm3 | yes |
| GAF | 2651 | GSE23466 | dm3 | yes |
| JIL-1 | 3035 | GSE27754 | dm3 | yes |
| mod(mdg4) | 324 | GSE20802 | dm3 | yes |
| Su(Hw) | 951 | GSE20833 | dm3 | yes |
| ZW5 | 3064 | GSE25373 | dm3 | yes |
| Fs(1)h | Pherson et al (2019) | GSE118484 | dm3 | yes |
| NippedB | Pherson et al (2019) | GSE118484 | dm3 | yes |
| Rad21 | Pherson et al (2019) | GSE118484 | dm3 | yes |
| SA | Pherson et al (2019) | GSE118484 | dm3 | yes |
| Smc1 | Pherson et al (2019) | GSE118484 | dm3 | yes |

**Table S7:** *Datasets for transcription and replication used in this work*

| Transcription and replication | | | dm3 or dm6 | LiftOver to dm6 |
|---|---|---|---|---|
| Orc2 | 2754 | GSE20888 | dm3 | yes |
| Topo-II | 5058 | GSE45069 | dm3 | yes |
| Pof | 3052 | GSE27808 | dm3 | yes |
| Pol-II | 950 | GSE20832 | dm3 | yes |
| 3'NT-seq | Pherson et al (2017) | GSE100545 | dm3 | yes |
| MED1 | Pherson et al (2019) | GSE118484 | dm3 | yes |
| MED30 | Pherson et al (2019) | GSE118484 | dm3 | yes |

**Table S8:** *Datasets for DNA accessibility used in this work*

| DNA accessibility | | | dm3 or dm6 | LiftOver to dm6 |
|---|---|---|---|---|
| DNase-I | Kharchenko et al (2011) | - | dm3 | yes |
| H1 | 3299 | GSE32767 | dm3 | yes |
| H2Av | 6073 | GSE45110 | dm3 | yes |
| H3 | 3302 | GSE32769 | dm3 | yes |
| H4 | 3303 | GSE32770 | dm3 | yes |

**Table S9:** *Datasets for histone modifications used in this work*

| Histone modificcations | | | dm3 or dm6 | LiftOver to dm6 |
|---|---|---|---|---|
| H2Bubi | 288 | GSE20771 | dm3 | yes |
| H3K18ac | 291 | GSE20774 | dm3 | yes |
| H3K23ac | 293 | GSE20776 | dm3 | yes |
| H3K27ac | 295 | GSE20778 | dm3 | yes |
| H3K27me1 | 3941 | GSE51965 | dm3 | yes |
| H3K27me2 | 2999 | GSE27789 | dm3 | yes |
| H3K27me3 | 297 | GSE20780 | dm3 | yes |
| H3K36me1 | 299 | GSE20782 | dm3 | yes |
| H3K36me3 | 301 | GSE20783 | dm3 | yes |
| H3K4me1 | 2653 | GSE23468 | dm3 | yes |
| H3K4me2 | 2654 | GSE23469 | dm3 | yes |
| H3K4me3 | 967 | GSE20839 | dm3 | yes |
| H3K79me1 | 3005 | GSE32736 | dm3 | yes |
| H3K79me2 | 306 | GSE20788 | dm3 | yes |
| H3K79me3 | 4934 | GSE45062 | dm3 | yes |
| H3K9me2 | 310 | GSE20791 | dm3 | yes |
| H3K9me3 | 312 | GSE20793 | dm3 | yes |
| H4K16ac | 316 | GSE20795 | dm3 | yes |
| H4K20me1 | 3286 | GSE32755 | dm3 | yes |
| H4K8ac | 5060 | GSE45070 | dm3 | yes |

**Table S10:** *Datasets for nucleosome remodelling factors used in this work*

| Nucleosome remodelling factors | | | dm3 or dm6 | LiftOver to dm6 |
|---|---|---|---|---|
| ASH-1 | 3279 | GSE32748 | dm3 | yes |
| JHDM1 | 5145 | GSE45092 | dm3 | yes |
| MRG15 | 3045 | GSE25365 | dm3 | yes |
| NURF301 | 5063 | GSE45072 | dm3 | yes |
| PR-Set7 | 5065 | GSE45074 | dm3 | yes |
| RPD3 | 4188 | GSE44523 | dm3 | yes |
| WDS | 5148 | GSE45094 | dm3 | yes |
| ISWI | 3030 | GSE27750 | dm3 | yes |
| MOF | 3041 | GSE27803 | dm3 | yes |

**Table S11:** *Datasets for polycomb and heterochromatin used in this work*

| Polycomb and heterochromatin | | | dm3 or dm6 | LiftOver to dm6 |
|---|---|---|---|---|
| Pc | 325 | GSE20803 | dm3 | yes |
| dRING | 927 | GSE20817 | dm3 | yes |
| sSFMBT | 2986 | GSE27728 | dm3 | yes |
| Ez | 2650 | GSE23465 | dm3 | yes |
| Pcl | 948 | GSE20830 | dm3 | yes |
| Psc | 3055 | GSE25370 | dm3 | yes |
| HP1a | 4126 | GSE44515 | dm3 | yes |
| HP1b | 3016 | GSE44462 | dm3 | yes |
| HP1c | 942 | GSE20824 | dm3 | yes |
| HP2 | 3026 | GSE27747 | dm3 | yes |
| HP4 | 4185 | GSE44521 | dm3 | yes |
| Su(var)3-7 | 2671 | GSE23486 | dm3 | yes |
| Su(var)3-9 | 952 | GSE20834 | dm3 | yes |

### Supplementary Materials and Methods

#### WT vs Mutant Borders Comparison

We called strong TAD borders using a value of threshold of the difference between the TAD-separation score of 0.08 and weak TAD borders using TAD-separation score of 0.04 excluding the borders that have been already annotated as strong. We compared TAD border positions between Wild Type (WT) and mutant and defined 10 types of changes:

- *strong -> strong* Border in WT has exactly the same position in mutant and annotated as strong in both WT and mutant.
- *weak -> weak* Border in WT has exactly the same position in mutant and annotated as weak in both WT and mutant.
- *strong -> weak* Strong border in WT has exactly the same position in mutant but became weak.
- *weak -> strong* Weak border in WT has exactly the same position in mutant but became strong.
- *strong -> fuzzy Strong b*order in WT is found within 2Kb window in mutant.
- *weak -> fuzzy* Weak border in WT is found within 2Kb window in mutant.
- *strong -> no* Strong border in WT is not found within 2Kb window in mutant.
- *weak -> no* Weak border in WT is not found within 2Kb window in mutant.
- *no -> strong* Strong border in mutant that does not belong to any class above.
- *no -> weak* Weak border in Mutant that does not belong to any class above.

We focused further analysis on two classes of borders. The borders that belong to *strong -> strong*, class we defined as *maintained borders*, and *strong -> no*, class we defined as *lost borders*.

#### Occupancy Analysis

For each class of borders (maintained and lost borders) we performed several preprocessing steps. First, we extracted the ChIP-chip signal within 5Kb window around WT TAD borders. Second, all signals within 5Kb window were winsorised: extreme values were replaced by cut-off points to reduce the effect of extreme outliers. For each single ChIP-chip dataset we selected 5%-quantile of negative signals as down cut-off point and 95%-quantile of positive signals as up cut-off point. Then, in order to have symmetric rescaling at the next step, we replaced the high cut-off point with the maximum of absolute values of cut-off points. The low cut-off was replaced by minus new up cut-off point value, respectively. After winsorisation, we divided all positive signals by up cut-off point and all negative signals by minus down cut-off point. The signals of resulting datasets belong to the [-1;1] interval that made them suitable for comparison. Note, that 3'NT-seq signal was analogically winsorised and rescaled. Also note that due to the high intensity of extreme signals, DNase-seq signal was winsorised with respect to zero down cut-off point and 75%-quantile up cut-off point. Then,

DNase-seq signal was rescaled by dividing by the maximum value. Third, we reordered borders with respect to the BEAF-32 signal summarised over 5Kb window.

**Bidirectional Transcription**

The directionality score computed as $\log_{10}$ of the ratio between nascent RNA levels in 500 bp on the positive strand downstream of the border and on the negative strand upstream of the border. 500bp bins that were 500bp away were considered in both directions from the border. The directionality score was sorted with respect to BEAF-32 summarised signal similar to ChIP-chip datasets. The borders with directionality score between -0.5 and 0.5 were treated as bidirectional and were coloured in green. Non-transcribed, positively transcribed and negatively transcribed borders were coloured in white.

**Clustering Analysis**

We performed clustering analysis on the maintained and lost borders. First, we extracted all ChIP-chip signals summarised over 5Kb window for each TAD border across all datasets and defined 50%-quantile of positive summarised signals as a limitQ (limitQ = 7.7041). Next, the limitQ was used for annotating each subset between six classes according to the following rules:

- *no signal* or *extra low signal* If the median of the subset summarised signals were less the zero. If 75%-quantile was greater than 0, then the subset was annotated as *extra low signal*, otherwise as *no signal*.
- *low signal* or *medium signal* If the median of the subset summarised signals were greater than the zero but less than limitQ. If 75%-quantile was greater than limitQ, then the subset was annotated as *medium signal*, otherwise as *low signal*.
- *high signal* or *extra high signal* If the median of the subset summarised signals were greater than limitQ. If 25%-quantile was greater than limitQ, then the subset was annotated as *extra high signal*, otherwise as *high signal*.
